# Temporal genomic analysis of melanoma rejection identifies regulators of tumor immune evasion

**DOI:** 10.1101/2023.11.29.569032

**Authors:** Sapir Cohen Shvefel, Joy A. Pai, Yingying Cao, Lipika R. Pal, Ronen Levy, Winnie Yao, Kuoyuan Cheng, Marie Zemanek, Osnat Bartok, Chen Weller, Yajie Yin, Peter P. Du, Elizabeta Yakubovich, Irit Orr, Shifra Ben-Dor, Roni Oren, Liat Fellus-Alyagor, Ofra Golani, Inna Goliand, Dean Ranmar, Ilya Savchenko, Nadav Ketrarou, Alejandro A. Schäffer, Eytan Ruppin, Ansuman T. Satpathy, Yardena Samuels

**Author notes:** Co-first authors (Sapir Cohen Shvefel, Joy A. Pai).

## Abstract

Decreased intra-tumor heterogeneity (ITH) correlates with increased patient survival and immunotherapy response. However, even highly homogenous tumors may display variability in their aggressiveness, and how immunologic-factors impinge on their aggressiveness remains understudied. Here we studied the mechanisms responsible for the immune-escape of murine tumors with low ITH. We compared the temporal growth of homogeneous, genetically-similar single-cell clones that are rejected vs. those that are not- rejected after transplantation *in-vivo* using single-cell RNA sequencing and immunophenotyping. Non-rejected clones showed high infiltration of tumor-associated-macrophages (TAMs), lower T-cell infiltration, and increased T-cell exhaustion compared to rejected clones. Comparative analysis of rejection-associated gene expression programs, combined with *in-vivo* CRISPR knockout screens of candidate mediators, identified *Mif* (macrophage migration inhibitory factor) as a regulator of immune rejection. *Mif* knockout led to smaller tumors and reversed non-rejection-associated immune composition, particularly, leading to the reduction of immunosuppressive macrophage infiltration. Finally, we validated these results in melanoma patient data.

**Statement of significance:** Here, we uncover the association of *Mif* expression with tumor growth and aggressiveness, specifically in low ITH tumors. These findings could facilitate the development of new strategies to treat patients with homogeneous, high-*MIF* expressing tumors that are unresponsive to immune checkpoint therapy.

## Introduction

Tumors with low intra-tumor heterogeneity (ITH), as determined by the number of clones comprising the tumor and the level of genomic clonal divergence, have been associated with improved survival and better responses to immunotherapy (1,2). Some factors underlying this correlation are known, but key aspects that explain the properties of ITH remain undetermined (3–5). Furthermore, accumulating evidence indicates that patients bearing low ITH tumors who suffer from aggressive disease show large variability in survival rates, suggesting that factors beyond ITH dictate disease course and tumor aggressiveness (6–8).

To elucidate the mechanisms that underlie clinical correlations and study the impact of ITH on immune-mediated tumor growth, we established a novel experimental system. This system builds on an *in vivo* mouse model, from Wolf et al., and allowed us to assess the effect of ITH independent of tumor mutational burden (5). Using this model, we hypothesized that we could investigate the mechanism(s) responsible for aggressive tumor behavior in low ITH patients. Indeed, we were able to compare tumors that are rejected and non-rejected while controlling for low ITH levels and differences in somatic variations.

We utilized this model system to comprehensively compare the transcriptomic state of tumor and immune cells from rejected and non-rejected tumors across several sequential time points during tumor evolution. Notably, we found that aggressive tumors showed specific immune phenotypes, including tumor-associated macrophage (TAM) infiltration, that adversely affected anti-tumor responses. Immune cell populations infiltrating the tumor microenvironment (TME) have been shown to have pro-tumorigenic effects by modulating T cell proliferation and cytotoxicity (9–11). Indeed, TAMs are known for their many contributions to such immune suppression, including direct inhibition of the cytotoxic T cell response, neoangiogenesis, secretion of pro-tumorigenic cytokines, and induction of the epithelial-mesenchymal transition (12–15). In addition, TAM infiltration is associated with poor prognosis and tumor progression in several cancer types including melanoma, pancreatic, breast, and bladder cancers (16–20), in line with our *in vivo* mouse data.

Analysis of gene expression time series data revealed *Mif* expression as a central non-genetic factor that affected the TME and dictated tumor growth in mice, which was subsequently validated in patient data. Indeed, previous reports have associated *Mif* upregulation with poor prognosis (21–25). Yet, its contribution to tumor aggressiveness is understudied. Our unique experimental system that isolated ITH from its complex context allowed the evaluation of MIF’s role in shaping the suppressive TME in a homogenous, low ITH tumors *in vivo*. Furthermore, our analysis suggests a means of potentially improving immunotherapeutic outcomes for patients bearing low ITH tumors.

## Results

### Survival time is variable in patients with low levels of ITH

Previous studies have suggested that melanoma patients with low ITH exhibit improved survival compared to patients with high ITH (1,2). However, analyzing the survival time of melanoma patients in The Cancer Genome Atlas (TCGA) partitioned according to high/low ITH levels revealed higher complexity. Although we found that the low ITH patients have significantly higher survival (**Supplementary Figure 1**A, Wilcoxon’s test, p=0.014) compared to high ITH, the distribution of survival times in low ITH patients is quite wide (**Supplementary Figure 1**A, interquartile range of 2,587 days for low ITH patients vs 1,452 days for high ITH patients). A different quantification of the TCGA survival data showed that 40% of the low ITH patients exhibit a poor outcome, with lower survival time than high ITH patients. These findings indicate that there is considerable variance in the response of low ITH patients (**Supplementary Figure 1**A). In addition, low ITH patients showed significantly higher survival time when characterized by a high T cell cytolytic scores (**Supplementary Figure 1**B). Taken together, these data motivated us to study factors that could contribute to immune and survival variability.

### An *in vivo* mouse model to identify factors that influence tumor aggressiveness in a low ITH setting

To examine factors that may play a role in tumor escape from immune surveillance, we established an experimental *in vivo* mouse model that would allow us to identify novel factors that contribute to tumor aggressiveness in low ITH tumors. To this end, we utilized the mouse model previously established in Wolf. et al, in which B2905, a mouse melanoma cell line (26), was exposed to ultraviolet B (UVB) radiation to increase ITH (5). From this heterogeneous parental line, we generated 40 homogeneous single-cell-derived clones (SCCs). SCCs were subjected to whole exome sequencing (WES), and analysis of the mutation variant allele frequencies (VAFs) verified that the clones harbored a narrow clonal VAF distribution with the fraction of clonal SNVs ranging from 0.835 to 0.949 (**Supplementary Table 1**).

Using this data we performed a phylogenetic analysis of the heterogeneous UVB cell line, which yielded a phylogenetic tree with eight terminal branches (TBs), numbered TB-1 to TB-8 (**Figure 1A**). We then placed the 40 SCCs on the various terminal branches of the tree, based on their sequence similarity. TBs were represented by clusters of 2 to 11 SCCs (**Figure 1A**). We focused on TB-3, represented by six SCCs that shared clonal and sub-clonal mutations, with an overall genetic similarity above 80% (**Supplementary Figure 1**C). Nevertheless, when inoculated into immunocompetent C57BL/6 mice, these SCCs exhibited very different growth phenotypes. Although five out of the six SCCs showed robust growth; two were rejected (SCC 31 and SCC 40) within 16 to 20 days post inoculation, while three were non-rejected (SCC 32, SCC 35, and SCC 37), growing aggressively and able to escape immune surveillance (**Figure 1B**). VAF values of the rejected and non-rejected clones were similar, demonstrating that the reason for the opposite growth phenotypes was not clonality (**Figure 1C**). To test whether remission of the rejected SCCs was dependent on the immune system, tumor volume was monitored after inoculation of rejected clones (SCC 31 and SCC 40) into NOD SCID gamma (NSG) immunodeficient mice. In the absence of a fully functioning immune system, all SCCs were able to grow and form tumors (**Figure 1D**), demonstrating that tumor rejection was indeed mediated by the immune system. Assessment of the MHC class I and II surface levels using flow cytometry validated that the aggressive phenotype was also not due to MHC class I downregulation (**Supplementary Figure 1**D). Analysis of WES data identified a single missense mutation in the gene Or13a22 (S5N) shared between the two rejected SCCs. No mutations were shared between the non-rejected SCCs (**Supplementary Figure 1**E, **Supplementary Table 2**), suggesting that any differences between rejected and non-rejected clones were likely due to non-genetic mechanisms, including first and foremost differentially expressed genes, and pathways in the tumor cells themselves, or in the TME.

**Figure 1.**
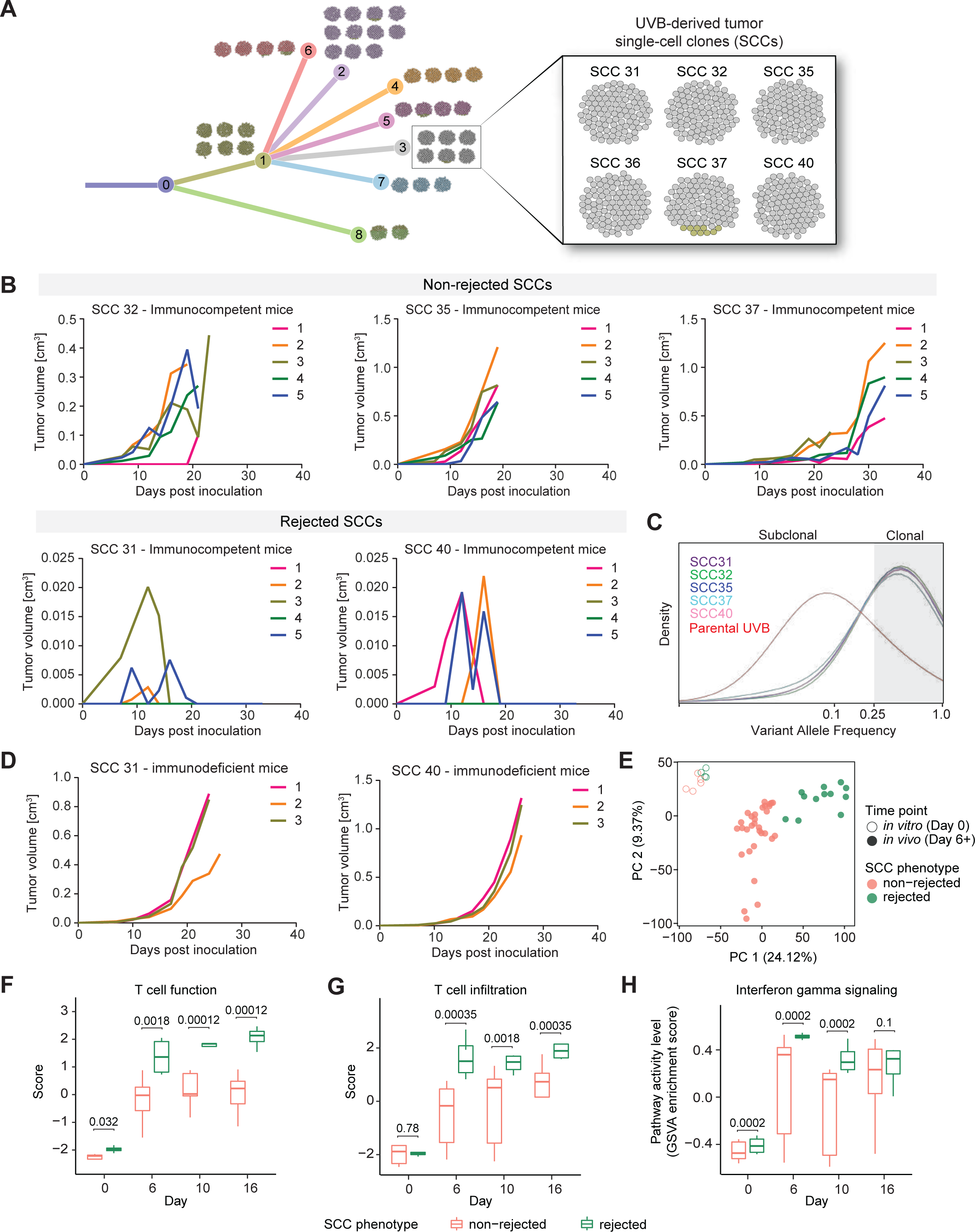
Single-cell clones with the same clonality and greater than 80% genetic similarity show opposite growth phenotypes. A) Phylogenetic tree representation of the UVB irradiated B2905 cell line. The tree depicts the results from mutation clustering analysis, which was used to define the distinct subclones present within the UVB cell line. The phylogenetic relationship between subclones is shown, and then each of the 40 UVB-derived SCCs is mapped onto the subclonal branch with the highest genetic similarity. Each of the 40 SCCs is depicted as a ball of 100 tumor cells, with the color coding reflecting the percentage frequency of each branch in each SCC sample. Shown in the top-left boxes is a representation of the UVB sample (median and mean VAF), again shown as a ball of 100 tumor cells, color-coded to match the subclonal branches. **B)** Growth curve of non-rejected (top) and rejected (bottom) SCC-derived tumors *in vivo* in immunocompetent mice. n=5. **C)** Distribution of variant allele frequencies (VAFs) of parental UVB-irradiated B2905 cells (red), rejected SCC 31 (purple) and SCC 40 (pink), and non-rejected SCC 32 (green), SCC 35 (blue) and SCC 37 (light-blue) in log2 space. VAF > 0.25 (log2=-2) is considered clonal. **D)** Growth curve of rejected SCC-derived tumors *in vivo* in NSG immunodeficient mice. n=3. **E)** A PCA plot based on the TMM-normalized log-CPM gene expression of the rejected (green) and non-rejected (pink) clones across *in vitro* (day 0; circles) and *in vivo* (day 6, 10, 16, 20; dots) time points after removing outlier samples (Methods). Top two principal components are shown, with the percentage of total variance explained labeled on the corresponding axis. **F)** T cell function scores, and **G)** T cell infiltration scores, both computed with the TIDE algorithm in the rejected (green) and non-rejected (pink) clones across different time points (X-axis). The scores between the rejected and non-rejected clones were compared with a linear model at each time point (Methods), and the corresponding BH-adjusted P values were shown. **H)** GSVA enrichment scores of the interferon-gamma pathway (Methods) in the rejected (green) and non- rejected (pink) clones across different time points (X-axis). Enrichment for the interferon-gamma pathway genes was tested with GSEA (Methods), and the corresponding BH-adjusted P values are shown.

### High TAM infiltration, low T cell frequency, and increased T cell exhaustion are associated with more aggressive tumor growth in low ITH tumors

To identify immune-specific cellular and transcriptional variation that may underlie the differential growth aggressiveness between highly clonal tumors, each SCC was inoculated separately into immunocompetent mice and the tumors were harvested at sequential time points (days 6, 10, and 16 post-inoculation for both groups, as well as day 20 post-inoculation for the non-rejected group). We then extracted and dissociated the tumors, and performed bulk RNA sequencing on these tumors, as well as on samples of the SCC cell lines representing day 0. Gene expression levels were compared between the SCC groups to identify differentially expressed genes and enriched pathways. PCA analysis revealed three well-defined clusters of expression patterns, corresponding to (a) samples from SCCs grown *in vitro* from day 0, (b) samples from the *in vivo* tumors of the non-rejected clones, and (c) samples from the *in vivo* tumors of the rejected clones, together establishing that the gene expression differences are only evident *in vivo* (**Figure 1E**). When comparing these groups, we found that the rejected tumors *in vivo* showed significantly increased T cell function and T cell infiltration signature scores compared to the non-rejected tumors (**Figure 1F, G**), as computed using the previously described Tumor Immune Dysfunction and Exclusion (TIDE) computational framework (27). In addition, we performed a gene set enrichment analysis on the differentially expressed sets of genes between the rejected and non-rejected groups (**Supplementary Tables 3, 4**). The expression of genes encoding IFNγ signaling pathway components, including *Camk2a*, *Irf1*, *Ifngr*, *Stat1*, *Jak1*, and *Jak2*, was consistently enriched in the rejected clones, both in the *in vitro* and *in vivo* time points (**Figure 1H**). IFNγ plays a central role in anti-tumor immunity by upregulating MHC-I expression and inducing the immunoproteasome (28). This pathway affects the response of melanoma patients and may thereby contribute to the observed rejection phenotype.

To fully decipher the differences in the tumor microenvironment of the rejected and non-rejected SCCs, we next performed single-cell RNA sequencing (scRNA-seq) on tumors harvested from mice transplanted with these SCCs at the same time points (days 0, 6, 10, 16, and 20 post-inoculation). Integration of cells from all tumors and time points revealed signatures of several major cell types, including a dominant population of tumor and stromal cells expressing Vimentin (*Vim*), *Cd81*, and *Pcbp2* (**Figure 2A**, **Supplementary Figure 2**A). We were also able to capture the immune compartment, which included adaptive immune cells such as T cells (*Cd3e, Cd3g)* and natural killer (NK) cells (*Nkg7*, *Gzma*), along with myeloid populations of monocytes (*Cd14*, *Fcgr1*), macrophages (*Cd74*, *Apoe*), and dendritic cells (DCs; *H2-Eb1*, *Il1b*) (**Supplementary Figure 2**A).

**Figure 2.**
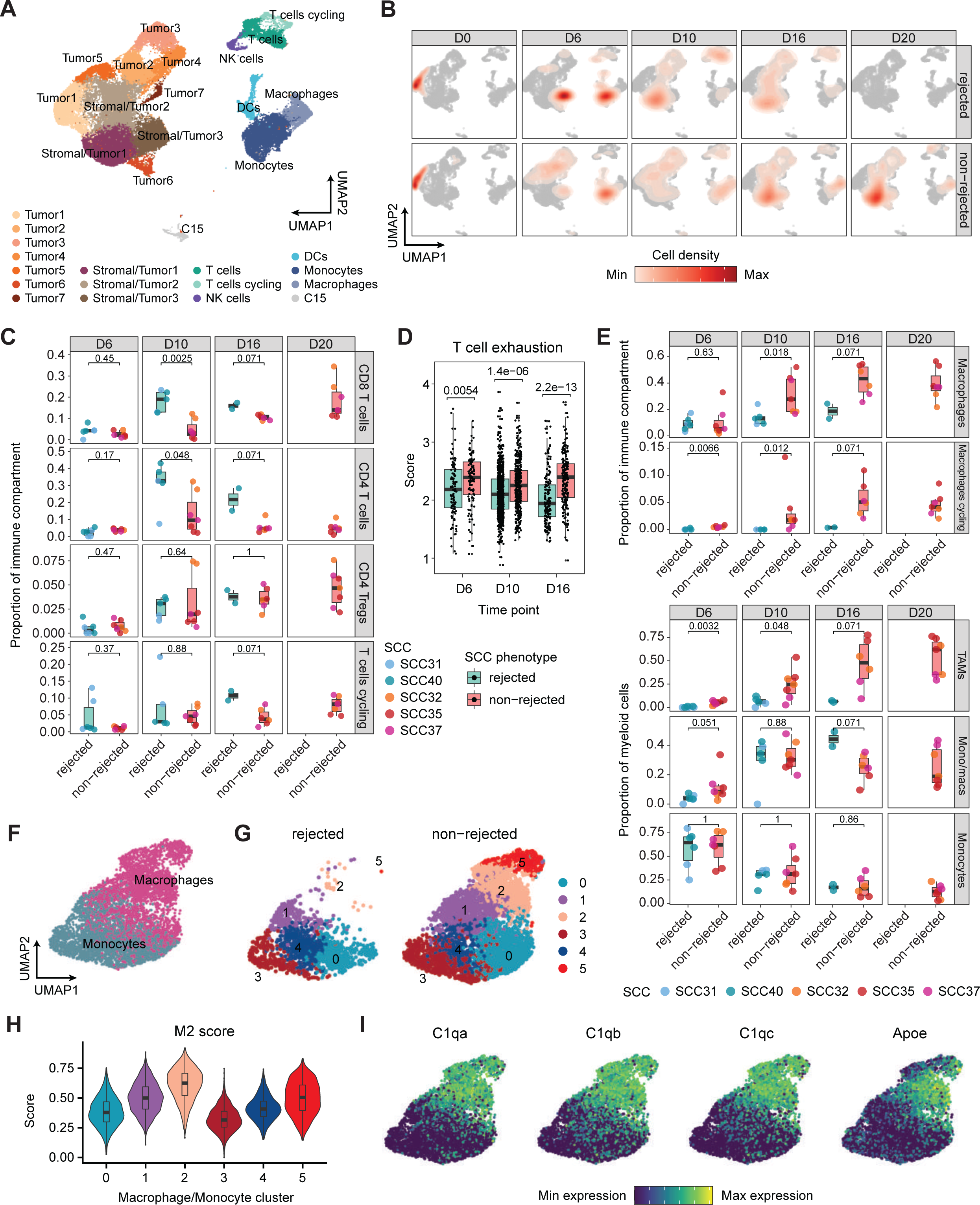
Single-cell RNA sequencing reveals increased M2-like macrophage infiltration in non- rejected tumors. A) UMAP of cell types obtained by scRNA-seq of rejected and non-rejected tumors. **B)** UMAP as in (A) overlayed with cell density per SCC group (rejected, non-rejected) and timepoint after tumor inoculation. **C)** Boxplot of cell type proportions of T cell clusters within the immune compartment of rejected and non-rejected SCCs at day 6, 10, 16, and 20 after tumor inoculation. Statistical testing by two-sided Wilcoxon test. **D)** Boxplot of CD8+ T cell exhaustion (84) scores between rejected (green) and non-rejected (red) tumors generated from SCCs from day 6 to day 16. The individual points denote the T cell exhaustion scores of CD8^+^ T cells across different replicates of a particular clone. P-values were determined using the Wilcoxon’s test between rejected and non-rejected at different time points. **E)** Boxplot of cell type proportions of macrophage clusters within the immune compartment (top) or myeloid compartment (bottom) of rejected and non-rejected SCCs at day 6, 10, 16, and 20 after tumor inoculation. Statistical testing by two-sided Wilcoxon test. **F-H)** Distribution of Mon/Mac subclusters between rejected and non-rejected groups, with M2 scores for each subcluster. **I)** Gene expression patterns of *C1qa*, *C1qb*, *C1qc*, and *Apoe* across different subclusters of Mon/Mac.

By comparing the rejected and non-rejected SCCs, we observed a higher proportion of macrophages in the non-rejected clones, especially at day 10 post-inoculation (**Figure 2B**, **Supplementary Figure 2**B). To further investigate the differential composition of the immune microenvironment between rejected and non-rejected tumors, we re-clustered cells in the immune compartment (**Supplementary Figure 2**C). This analysis enabled a finer resolution of the T cell compartment, separating T cells into cytotoxic CD8^+^ T cells (*Nkg7*, *Ccl5*), CD4^+^ regulatory T cells (*Foxp3, Ctla4*), conventional CD4^+^ T cells (*Ifng*, *Rora*), and proliferating T cells (*Mki67*) (**Supplementary Figure 2**D). Comparison of T cell populations from the non-rejected versus rejected tumors revealed a significantly decreased CD8^+^ and conventional CD4^+^ T cell frequency, and a higher exhaustion phenotype in the non-rejected tumors (**Figures 2C, D**). Furthermore, we separated macrophages into resting and proliferating populations based on the expression of *Mki67* (**Supplementary Figure 2**C**, E**). Both macrophage populations accounted for a significantly higher fraction of the immune compartment in non-rejected tumors compared to rejected tumors at day 10 post-inoculation (**Figure 2E**). Additionally, the proliferating macrophages were present at higher frequencies in the non-rejected tumors as early as day 6, suggesting that the TME of non-rejected SCCs induces the proliferation of macrophages shortly after tumor formation, which leads to the accumulation of macrophages in the tumor. A similar analysis of myeloid cells also showed an enrichment of macrophages in non-rejected SCCs at significant levels on days 6 and 10 post-inoculation (**Figure 2E**, **Supplementary Figure 2**F). When we specifically analyzed CD14^+^ monocytes and macrophages, we were able to identify six clusters, two of which (clusters 2 and 5) were found to be significantly enriched in the non-rejected tumors (**Figure 2F, G, Supplementary** Figure 2G). These two clusters were characterized by a high M2 pro-tumorigenic score (**Figure 2H**), as well as high expression of *C1qa, C1qb, C1qc,* and *Apoe* which are known to be associated with M2 macrophages (**Figure 2I**) (29). These data further strengthen our observation that the non-rejected tumors are associated with relatively high numbers of pro-tumorigenic macrophages compared to rejected SCC-derived tumors.

Opal multiplexed immunohistochemistry (IHC) staining of the rejected and non-rejected tumors ten days post-inoculation validated the scRNA-seq results, showing significantly higher CD206^+^CD204^+^ pro-tumorigenic macrophages and lower CD8^+^ T cell infiltration (**Figure 3A-E**). Importantly, a T cell suppression assay, which assesses T cell proliferation, established that co-culturing non-rejected SCC tumor-derived macrophages with CFSE-labeled CD8^+^ or CD4^+^ T cells derived from naïve mouse spleens inhibited T cells from proliferating. Furthermore, it significantly decreased their cytotoxic activity, manifested by significantly lower levels of IFNγ secretion (**Figure 3F**). This data corroborates the role of macrophages that infiltrate highly aggressive tumors in suppressing T cell function, ultimately leading to tumor immune escape.

**Figure 3.**
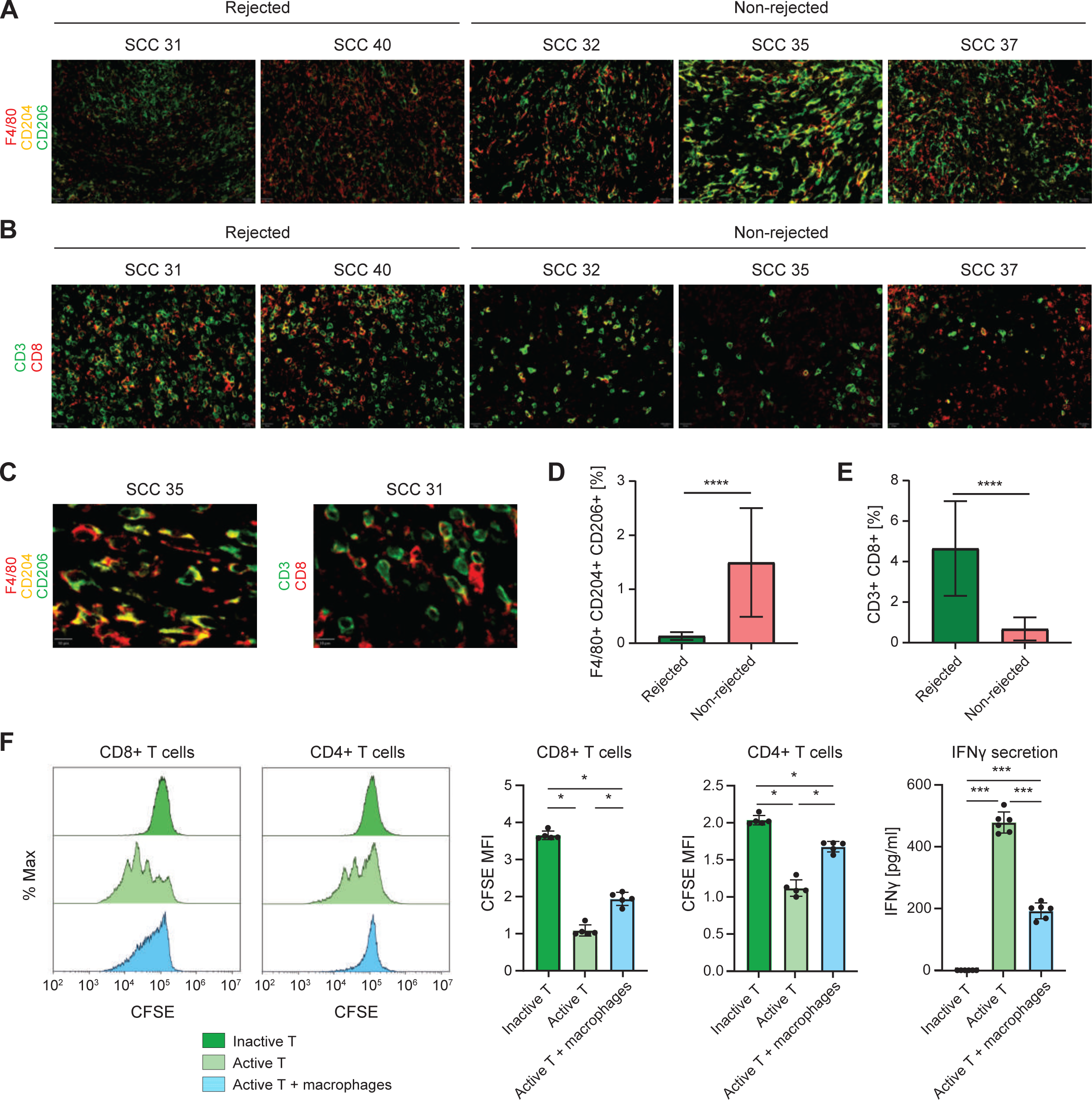
Opal staining data confirms increased M2-like macrophage infiltration in the non-rejected single-cell clones. A) Representative Opal Multiplex immunohistochemical stains for F4/80 (red), CD204 (yellow), and CD206 (green) in tumors derived from SCC 31, SCC 40, SCC 35, SCC 37, and SCC 32 on day 10 after inoculation. 4 areas from each tumor and 3 tumors from each single cell clone were examined. Scale bars represent 20 μM. DAPI staining is not shown. **B)** Representative Opal Multiplex IHC stains for CD3 (green) and CD8 (red) in tumors derived from SCC 31, SCC 40, SCC 35, SCC 37, and SCC 32 on day 10 after inoculation. 4 areas from each tumor and 3 tumors from each single cell clone were examined. Scale bars represent 20 μM. DAPI staining is not shown. **C)** Larger magnification of representative Opal Multiplex immunohistochemical stain for F4/80^+^CD204^+^CD206^+^ cells in tumor derived from SCC35 (left) and CD3^+^CD8^+^ cells in tumors derived from SCC 31 (right) on day 10 after inoculation, to demonstrate markers co-localization. Scale bars represent 10 μM. DAPI staining is not shown. **D)** Quantification of the percentage of F4/80+CD204+CD206+ cells described in (A). Data are mean ± SEM. The rejected and non-rejected groups were compared using Mann-Whitney U test with a p-value of 2.55*10^-8^. **E)** Quantification of the percentage of CD3+CD8+ cells described in (B). Data are mean ± SEM. The rejected and non-rejected groups were compared using Mann-Whitney U test with a p-value of 7.094*10^-7^. **F)** CFSE-based T cell proliferation assay following 48 hours co-culture with non-rejected SCC35 tumor-derived macrophages isolated 10 days post-inoculation. Representative histogram plot and quantification of CFSE intensity (n = 5). CD8^+^/CD4^+^ T cells isolated from healthy mouse spleens cultured w/o macrophages served as control. CD8: The Kruskal-Wallis (KS) chi-squared test shows Chi-square(2) 12.5, P = 0.001930454. Pairwise comparison by Wilcoxon test with Bonferroni correction are shown in the figure. * Pvalue < 0.05. CD4: The Kruskal-Wallis (KS) chi-squared test shows Chi-square(2) 12.5, P = 0.00193. Pairwise comparison by Wilcoxon test with Bonferroni correction are shown in the figure. * Pvalue < 0.05. T cell mediated IFNγ secretion to the culture media, 48 hours post co-culture of T cells with or without tumor-derived macrophages, measured using an ELISA. n=6. Data are mean ± SEM. The Kruskal-Wallis (KS) chi-squared test shows Chi-square(2) 15.726, P = 0.0003847. Pairwise comparison by Wilcoxon test with Bonferroni correction are shown in the figure. *** Pvalue < 0.01.

### Increased *Mif* levels lead to aggressive tumor growth of low ITH tumors

We next sought to identify tumor-intrinsic factors that may contribute to the dramatic differences seen in the immune cell composition of non-rejected (derived from SCC 32, SCC 35, and SCC 37) vs. rejected (derived from SCC 31 and SCC 40) tumors. Although all tumor populations shared a core identity, reflected in expression of specific genes such as *Vim*, *Gpx4*, and *Pcbp2* (**Supplementary Figure 2**A), we identified numerous transcriptional differences between the aggressive and non-aggressive tumors. First, we performed differential expression analysis between the tumor compartment of non-rejected and rejected clones at each time point (0, 6, 10, and 16 days post-inoculation) separately. In total, we observed 128 differential genes at day 0 (101 up in non-rejected, 27 up in rejected), 282 genes at day 6 (123 non-rejected, 159 rejected), 169 genes at day 10 (79 non-rejected, 90 rejected), and 204 genes at day 16 (100 non-rejected, 104 rejected; fold change >1.5; Bonferroni-corrected p-value <0.01, Wilcoxon rank sum test; **Supplementary Table 5, Methods**). To investigate whether there are shared transcriptional programs between individual clones that control tumor aggressiveness, we identified a core set of genes that were upregulated in either all rejected or all non-rejected clones for each time point. This revealed 4, 11, and 13 core genes that were overexpressed in both rejected clones at days 0, 6, and 10, respectively (**Supplementary Figure 3**A). Additionally, we identified 29, 13, and 11 core non-rejection genes at days 0, 6, and 10, respectively (**Supplementary Figure 3**B). Of note, we observed that at day 0, all non-rejected clones expressed higher levels of calreticulin (*Calr*), protein isomerase *Pdia3*, and members of the S100 protein family (*S100a4*, *S100a6*), which have been associated with tumor progression and increased metastasis (30). Next, we identified genes that were consistently differentially expressed between rejected and non-rejected SCCs across time points. This analysis revealed 78 genes that were upregulated by rejected clones at two or more time points, including the arginine metabolism enzyme *Ass1*, whose expression has been linked to tumor suppression and better prognosis of patients with breast cancer and hepatocellular carcinoma (31) (**Figure 4A**). Among the non-rejected tumor clones, we found 91 genes to be upregulated at two or more time points, including Macrophage migration inhibitory factor (*Mif*), *Lgals1*, and *S100a11* (**Figure 4A**). Since these genes displayed consistent upregulation in non-rejected SCCs longitudinally, we hypothesized that they could be factors that mediate tumor aggressiveness.

**Figure 4.**
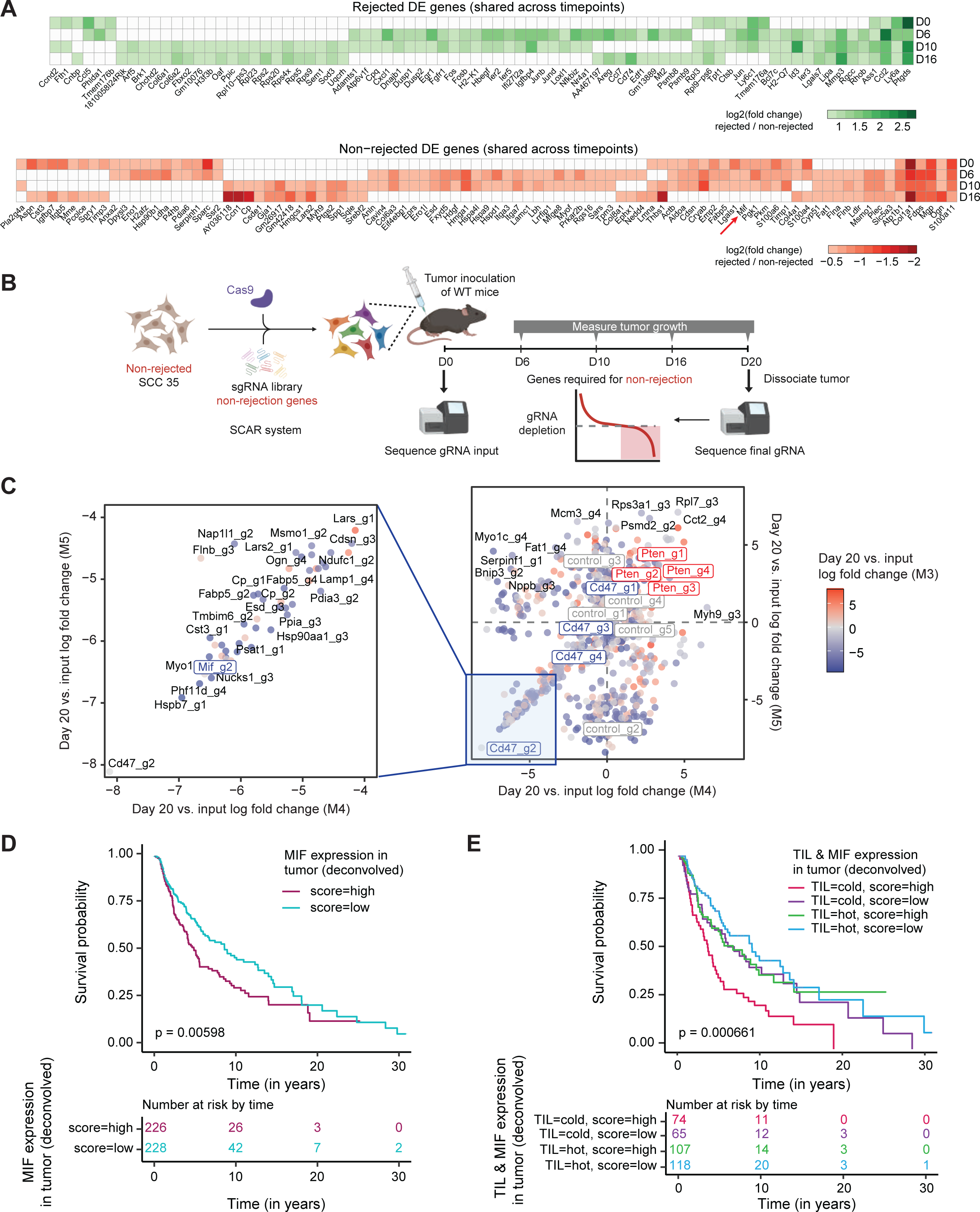
*Mif* is overexpressed in non-rejected clones and in melanoma patients with poor survival. **A)** Differentially expressed genes in the rejected (top) and in the non-rejected (bottom) group of clones that are recurrent across several time points, identified in the scRNA-seq analysis. *Mif* is shown to be differentially expressed in the non-rejected clones in 3 time points. **B)** Schematic of pooled *in vivo* CRISPR screen for non-rejection genes. **C)** sgRNA depletion and enrichment of CRISPR-edited libraries. The score represents the average log fold change of guide counts at day 20 post-inoculation and day 0 input libraries. Highlighted in the left square are the top depleted guides. **D)** Kaplan-Meier survival curves (time is measured in years on the x-axis) of de-convolved gene expression (37) of TCGA melanoma patients stratified by *MIF* expression in tumor cells. P-values were determined using the log-rank test to compare the overall survival probability between patients with high (in red) and low (in blue) *MIF* expression. **E)** Kaplan-Meier survival curves (time is measured in years on the x-axis) of de-convolved gene expression (37) of TCGA melanoma patients stratified by a combination of Tumor Infiltrated Lymphocyte (TIL) patterns (pathological annotation of tumors as having a hot or cold tumor microenvironment as previously described (38) and by *MIF* expression in tumor cells. P-values were determined using the log-rank test to compare the overall survival probability between patients with the combination of tumor hot/cold patterns with high/low *MIF* expressions.

To narrow down the list and rationally choose the best targets for further exploration, we performed an *in vivo* CRISPR screen, which allowed us to examine which genes may be required for immune evasion. To this end, we created a pooled CRISPR sgRNA library targeting 200 genes that were differentially expressed in the non-rejected tumors at day 0, 6, 10, or 16 post-inoculation. We also included 5 non-targeting control guides and a set of 3 depletion and 2 enrichment control genes, including *Cd47* and *Pten*, respectively, based on previous studies (32). We then transduced the non-rejected SCC 35 cell line with the sgRNA library and Cas9 using selective CRISPR antigen removal (SCAR) lentiviral vectors in order to induce per-cell knock-outs *in vitro,* while avoiding immune-mediated rejection of CRISPR/Cas9 components *in vivo* (**Figure 4B**) (33,34). The edited SCC35 cell lines were then split into 3 replicates at day 0 and sequenced to verify high sgRNA recovery rates, which ranged from 93.5%-95.6% of the original guide library (**Supplementary Figure 4**A). These input pools were then inoculated into immunocompetent mice, after which the tumors were harvested, dissociated, and subjected to sgRNA sequencing 20 days post-inoculation (**Figure 4B**, **Supplementary Figure 4**B). At day 20, we recovered between 40.6%-73.6% of sgRNAs in the pooled library, which was lower than in the input library at day 0, likely reflective of *in vivo* selection pressures (**Supplementary Figure 4**A). By comparing this final sgRNA library to the input library sequenced at day 0 and assessing guide enrichment/depletion using a negative binomial model (**Methods**), we identified 451 significantly depleted and 397 enriched guides, and 33 significantly depleted genes and 38 enriched genes (FDR-adjusted p-value <0.05, absolute log fold change >1). We first analyzed screen controls and found that all 4 guides targeting the enrichment control *Pten* were enriched at day 20 in both replicates M4 and M5 (4/4 guides significantly enriched at adjusted p-value <0.05 in M4; 2/4 guides significant in M5) (**Figure 4C**, **Supplementary Figure 4**C**, D**). Furthermore, the depletion control *Cd47* was depleted in all replicates at the guide level (4/4 guides depleted in M3; 4/4 in M4; 3/4 in M5) and gene level (adjusted p-value 0.000171 in M4), suggesting that the results of our pooled screen recapitulated known effects of gene knockouts on tumor aggressiveness. Indeed, the most strongly depleted guide in the entire CRISPR screen for replicates M4 and M5 was Cd47_g2 (M4: log fold change -8.12, FDR-adjusted p-value 0.02; M5: log fold change -8.11, FDR-adjusted p-value 0.02) (**Figure 4C**, **Supplementary Figure 4**C).

One of the next most strongly depleted hits was a guide targeting Macrophage migration inhibitory factor (*Mif*), (Mif_g2; M4: log fold change -6.31, FDR-adjusted p-value 0.05; M5: log fold change -6.24, FDR-adjusted p-value 0.05) (**Figure 4C**), a gene whose expression was also consistently enriched in non-rejected clones longitudinally from days 0 to 10 post-inoculation (**Figure 4A**). Additionally, *Mif* was significantly depleted at the gene level in replicate M4 (median log fold change -2.5, FDR-adjusted p-value 0.0099). MIF is a secreted protein that binds to the CD74 receptor, which is highly expressed on professional antigen-presenting cells such as macrophages and dendritic cells (35). Given that MIF is secreted by the tumor cells in the TME, we hypothesized that it may recruit CD74^+^ monocytes and promote their differentiation into pro-tumorigenic macrophages. This would then contribute to an immunosuppressive environment and cause tumor growth (36). This hypothesis is consistent with the increased infiltration of TAMs with a high M2 score and high expression levels of the *C1q* and *Apoe* genes in the non-rejected tumors observed by scRNA-seq analysis (**Figure 2E-I**).

To assess the relevance of our mouse findings to human melanoma, we computed the correlation between *MIF* expression and tumor aggressiveness in de-convolved-by-cell-type gene expression in TCGA melanoma patient datasets, which was generated via the application of CODEFACS to the TCGA bulk expression data (37). This analysis revealed significantly poorer survival of patients estimated to express high *MIF* levels in tumor cells (**Figure 4D**; log-rank test, p = 0.00598). Importantly, we also analyzed de- convolved gene expression data of tumors annotated as having a tumor-infiltrating lymphocyte (TIL) pattern (38) and found accentuated poor survival for patients bearing both cold tumors and estimated high *MIF* expression in tumor cells (**Figure 4E**; log-rank test, p = 0.000661). These results further highlight *MIF*/*Mif* activity in tumor cells as a possible factor that underlies immune escape by non-rejected clones.

### *Mif* KO in the aggressive clones significantly reduced tumor growth

To assess the contribution of *Mif* to the aggressive growth phenotype in mice, we knocked out (KO) *Mif* using the CRISPR/Cas9 system in both non-rejected SCC35 (KO clones 19 and 24) and SCC37 (KO clone 21) (**Supplementary Figure 5**A). Injection of these KO clones into immunocompetent mice gave rise to significantly smaller tumors compared to the parental *Mif*-wildtype (WT) or control cells (cells that were transfected with the same CRISPR/Cas9 vectors, sorted for GFP^+^ phenotype, grew as SCCs, but found to have WT *Mif* by Sanger sequencing) (Wilcoxon p-value 0.0001 at day 21 and 0.00004 at day 27, for SCC35 and SCC37, respectively) (**Figure 5A**, **Supplementary Figure 5**B).

**Figure 5.**
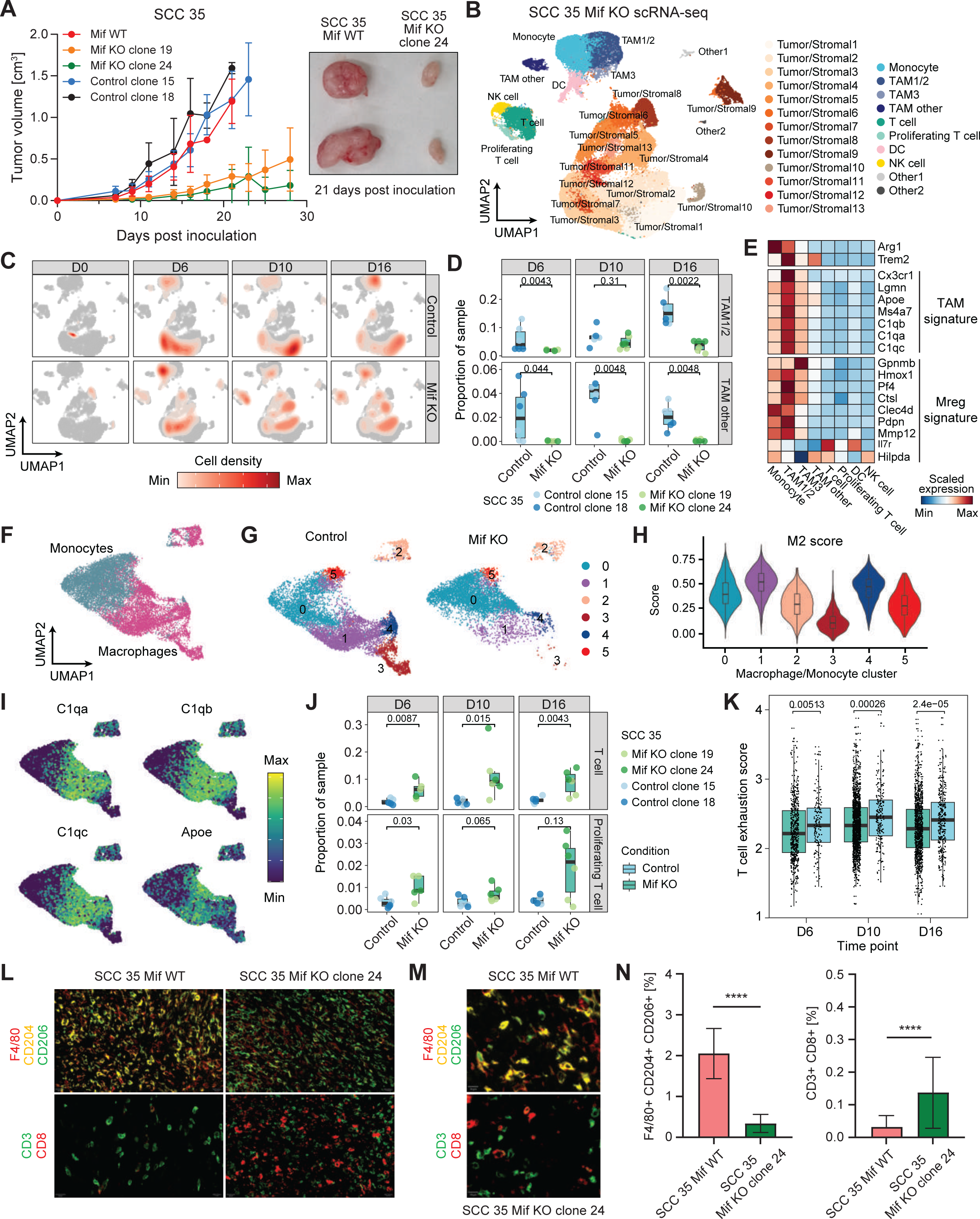
Decreased macrophage and increased T cell infiltration of SCC-derived tumors with *Mif* KO. A) *Mif* KO using CRISPR in the non-rejected SCC35 significantly decreased tumor growth compared to the *Mif* WT control clones. Statistical analysis between *Mif* KO to WT expressing clones at day 21 (the latest time point in which all groups are represented) gave a P value of 0.0001 by two-sided Wilcoxon-test followed by Bonferroni correction. n=8-10.**B)** UMAP of cell types obtained by scRNA-seq of SCC35- derived tumors with or without *Mif* KO. Data shows samples from all tumors and time points aggregated together. **C)** UMAP as in (B) overlayed with cell density per condition (Control or *Mif* KO) and timepoint after tumor injection. **D)** Boxplot of cell type proportions of macrophage clusters from control and *Mif* KO clones at day 6, 10, 16, and 20 after tumor inoculation. Statistical testing by two-sided Wilcoxon-test (* <0.05, ** <0.01, ***<0.001). **E)** Heatmap of tumor-associated macrophage markers among immune cell type clusters. **F-G)** Distribution of Mon/Mac subclusters between *Mif* KO and control groups. **H)** M2 score for each subcluster of Mon/Mac. **I)** Gene expression patterns of *C1qa*, *C1qb*, *C1qc*, and *Apoe* across different subclusters of Mon/Mac. **J)** Boxplot of cell type proportions of T cell clusters from control and *Mif* KO clones at day 6, 10, 16, and 20 after tumor inoculation. Statistical testing by two-sided Wilcoxon- test (* <0.05, ** <0.01, ***<0.001). **K)** Boxplot of general T cell exhaustion (84) scores between *Mif* KO (green) and control (blue) tumors generated from SCC35 from day 6 to day 16 (legend as in (J)). The individual points denote the T cell exhaustion scores of T cells across different replicates of a particular clone. P-values were determined using the Wilcoxon’s test between rejected and non-rejected at different time points. **L)** Representative Opal Multiplex IHC stains for: Top: F4/80 (red), CD204 (yellow), and CD206 (green), Bottom: CD3(green) and CD8 (red) in tumors derived from *Mif* WT or KO SCC35 on day 10 after inoculation. 4 areas from each tumor and 3 tumors from each single cell clone were examined. The scale bars represent 20 μM. DAPI staining is not shown. **M)** Larger magnification of representative Opal Multiplex immunohistochemical stain for F4/80^+^CD204^+^CD206^+^ cells in tumor derived from SCC35 *Mif* WT (top) and CD3^+^CD8^+^ cells in tumors derived from SCC 35 *Mif* KO (bottom) on day 10 after inoculation, to demonstrate markers co-localization. Scale bars represent 10 μM. DAPI staining is not shown. **N)** Quantification of the percentage of F4/80^+^CD204^+^CD206^+^ cells (left) or CD3^+^CD8^+^ cells (right) described in (L). Data are mean ± SEM. Mann-Whitney U test. P-values are 5.015*10^-6^ and 1.324*10^-6^, respectively.

We next assessed whether tumors from *Mif* KO SCCs exhibited changes in the tumor microenvironment compared to tumors established from the rejected SCCs. We performed scRNA-seq of tumors derived from SCC35-derived *Mif* KO clones as well as control SCC35 tumors at days 0, 6, 10, and 16 post-inoculation.

After integrating data from all time points and clones, we observed that the broad cell types recapitulated those from scRNA-seq of the rejected vs. non-rejected SCCs **(Figure 5B, 2A**). Namely, we captured a large population of tumor/stromal cells, while the immune compartment was composed of macrophages, monocytes, DCs, NK cells, and T cells in both resting and proliferating states (**Figure 5B**). We confirmed that *Mif* expression was indeed lower among tumor cells in *Mif* KO clones compared to control clones (**Supplementary Figure 5**C).

Next, we examined TAMs in *Mif* KO clone-derived tumors and observed decreased infiltration compared to the controls (average frequencies of TAM1/2 at day 6: 2.0% *Mif* KO, 5.9% control, Wilcoxon p-value 0.0043; day 10: 4.7% *Mif* KO, 6.8% control. Wilcoxon p-value 0.31; day 16: 3.5% *Mif* KO, 16.0% control, Wilcoxon p-value 0.0022) (**Figure 5B-D**). These TAMs were associated with M2 characteristics, exhibiting a high M2 score and *C1qa, C1qb, C1qc,* and *Apoe* expression (**Figure 5E-I, Supplementary** Figure 5D). Conversely, the myeloid population in the *Mif* KO clones was skewed towards the monocyte cluster (**Figure 5C**). The abundance of TAMs with a high M2 score and *C1qa, C1qb, C1qc,* and *Apoe* expression seen in macrophage clusters 1 and 4 (**Figure 5H**) in controls compared to *Mif* KO groups is consistent with our finding that TAMs with the highest M2 scores and high levels of expression of *C1qa, C1qb, C1qc,* and *Apoe* genes (cluster 2 and 5 in **Figure 2G-I**) are more abundant in non-rejected groups. We also found that *Mif* KO samples exhibited higher T cell infiltration compared to the controls (average frequencies of T cells at day 6: 7.3% *Mif* KO, 2.0% control; day 10: 12.3% *Mif* KO, 2.2% control; day 16: 10.7% *Mif* KO, 2.8% control) (**Figure 5J**). To further characterize the differences in T cell infiltration induced by the *Mif* KO, we re-clustered the T cell populations (**Supplementary Figure 5**E). This analysis enabled the separation of T cells into a naïve/memory cluster characterized by the expression of *Ccr7* and *Tcf7*, and two clusters of CD4^+^ T cells differentiated by *Foxp3* expression (**Supplementary Figure 5**E**, F**). We identified two clusters of CD8^+^ T cells expressing cytotoxicity markers including *Gzma*, *Gzmb*, and *Nkg7* that were separated based on the expression of the proliferation marker *Mki67* (**Supplementary Figure 5**F). Additionally, we captured a cluster of T cells that expressed high levels of interferon-stimulated genes such as *Ifit1* and *Isg15*. By comparing these populations between conditions, we observed a higher frequency of cytotoxic CD8^+^ T cells in *Mif* KO clones as early as day 6 post-inoculation (4.7-fold increase at day 6, 3.0- fold increase at day 10, 9.8-fold increase at day 16) (**Supplementary Figure 5**G). Furthermore, the frequency of CD8^+^ T cells increased over time in the *Mif* KO clones, whereas it stayed relatively constant in control samples (**Supplementary Figure 5**G).

We also observed a higher frequency of regulatory CD4^+^ T cells (Tregs) in the control by day 16 post-inoculation (**Supplementary Figure 5**G). These differences in T cell composition were accompanied by a lower T cell exhaustion signature in *Mif* KO clones (**Figure 5K**). To validate these findings, IHC staining was performed on tumor samples of the *Mif* WT or KO SCC35 10 days post-inoculation, revealing significantly decreased CD206^+^CD204^+^ macrophages and increased CD8^+^ T cells infiltration to the KO tumors (**Figure 5L-N**). Altogether, these data support our hypothesis that *Mif* expression and function plays a central role in inducing TAM infiltration followed by T cell dysfunction, both contributing to the immunosuppressive microenvironment and ultimate aggressive tumor growth.

## Discussion

Here we have established an experimental system to study genetically similar homogenous tumors and assess the factors that impact their aggressiveness in a controlled manner. We established 40 cell lines derived from 40 single-cell clones that were deeply annotated genetically, establishing their mutational landscape, clonality, and phylogenic relations. This approach provided the opportunity to perform a comparative analysis of rejection-associated gene expression signatures, accompanied by *in vivo* CRISPR knockout screens of promising candidate regulators of tumor aggressiveness, measuring their influence on anti-tumor immunity and tumor growth. Our findings in mice suggest that, in melanoma, an important determinant of an aggressive growth phenotype in low ITH tumors is *Mif* expression in tumor cells, whose activity is associated with increased TAM frequency and decreased cytotoxic T cell infiltration into TME.

The infiltrating TAMs are characterized by a pro-tumorigenic phenotype, manifested by a high M2 score and elevated expression of *C1qa*, *C1qb*, and *C1qc*, which encode subunits of the C1q component of the complement system, along with *Apoe* (29,39). These genes have been previously reported as overexpressed in TAMs in various tumor types, including melanoma, and are generally associated with immunosuppression and poor prognosis (40–42). Validation of this finding using IHC staining provided further information regarding the spatial location of these TAMs, which infiltrate the tumor core and comprise high levels of CD206^+^CD204^+^ macrophages. These results corroborate previous reports that such macrophages are frequently found in patients bearing aggressive tumors and are associated with poor survival in various cancers, including melanoma (43–48). Indeed, patient data analysis shows that higher TAM infiltration is associated with worse prognosis for various cancers, further supporting this observation (49). Furthermore, high TAM infiltration has been reported to be accompanied by increased CD8^+^ T cell exhaustion and increased presence of Tregs in the TME (29). Indeed, our scRNA-seq sequencing results revealed reduced T cell infiltration and function along with a high exhaustion phenotype.

We undertook a comprehensive genomic analysis to pinpoint mechanisms underlying these distinct immune microenvironment phenotypes. Employing temporal analysis of differentially expressed genes shared between the cancer cells of several non-rejected tumors, followed by CRISPR perturbation screen to narrow down the candidates, identified *Mif* as a key factor dictating tumor ability to evade immune surveillance.

Our results showed significantly and consistently higher Mif expression in the non-rejected tumors compared to the rejected tumors across several time points of the tumor evolution, starting from the *in vitro* day 0 samples. Unequivocal proof of our hypothesis that the aggressive growth phenotype is mediated by *Mif* came from the dramatic phenotype reversal upon *Mif* knockout in the non-rejected SCCs. Indeed, these *Mif* KO SCCs reverted their aggressive phenotype and showed an immune infiltration profile that was similar to the rejected tumors.

Our experimental mouse data are recapitulated in TCGA melanoma patients, where the overall survival rate is significantly higher in tumors expressing low *MIF* levels, a result which is even more pronounced when also considering T cell infiltration. These findings, further support the detrimental influence of high *MIF* levels on the anti-tumor immune response in humans, in keeping with previous studies (21–24). In the future, it will be important to evaluate whether the response of highly homogeneous tumors to ICB correlates with *MIF* expression levels. This analysis is currently impossible due to the low number of such patients.

We suggest that in the subset of highly homogeneous tumors that overexpress *Mif*; it binds to the CD74 receptor (35,36,50), potentially recruiting CD74^+^ macrophages. These CD74^+^ macrophages, differentiate into TAMs and exhibit an M2 signature. These M2 macrophages, in turn, reduce CD4^+^ and CD8^+^ T cell infiltration into the tumor. The total outcome is weaker anti-tumor immunity, manifested by dampening of TIL cytotoxicity, effector cytokine secretion, proliferation, and a high T cell exhaustion signature. In contrast, highly homogeneous tumors that harbor reduced *Mif* expression also have low presence of macrophages, and specifically those in a proliferative state. In addition, a significantly lower TAM presence in the tumor leads to enhanced CD4 and CD8 T cell infiltration into the tumor and elevated effector cytokines, enhancing the ability of the immune system to reject the tumor.

Although high *MIF* expression has previously been associated with a more robust immunosuppression phenotype, increased tumor growth, and poor outcome (21–25,51), its association with worse prognosis has not always been clear cut, as there have been cases where it was shown to be associated with better prognosis (52,53). Our focus in this study on highly homogeneous tumors, and our establishment of an experimental model that emphasizes the use of SCC-derived, highly clonal tumors, enabled us to identify *Mif* as a consistent and robust biomarker for tumor aggressiveness. Indeed, our *in vivo* CRISPR screen was based on a heterogeneous system where we assessed a pool of targets in parallel. This may explain our ability to identify *Mif* as a target. Our follow up, single *Mif* KO functional data, supports this hypothesis as in this case we observed a clear-cut phenotype. It may therefore be the case that, *Mif*, like many additional targets, was missed in previous CRISPR screens due to the tumors’ heterogeneous nature, as well as the fact that the impact of secreted factors is masked by other cells in the screen, and their results are diluted.

Our findings show the value in evaluating mechanisms of resistance in a highly homogeneous system. Importantly, beyond *Mif*, our unique experimental system identified additional genes enriched in aggressive tumors throughout several time points of the tumor’s evolution that have not been identified in former studies.

*MIF* could therefore serve as a potential target for anti-cancer treatments, specifically in low ITH cases (54). MIF inhibitors, such as ISO-1, ISO-66, and CPSI-1306, have been shown to decrease tumor growth in melanoma, and clinical trials of IPG1094, another MIF inhibitor, led to FDA approval for the treatment of several tumor types (22,55–62). Our highly homogeneous model can therefore serve as an effective platform to evaluate the effect of MIF inhibitors on tumor growth and immune cell infiltration in homogenous non-rejected SCCs. In conclusion, we suggest that MIF levels in cancer cells are a strong determinant of the immune response in highly homogenous melanoma tumors, highlighting the potential importance of assessing it as a target for future therapies.

## Author Contributions

Y.S., A.S, S.C.S and J.P designed the study, performed experiments, supervised the project and wrote the paper. J.P, Y.C, L.R, K.C and R.L analyzed the RNAseq, WES, and patient data. M.Z analyzed the IHC stained images. J.P and W.Y performed and analyzed the CRISPR screen. O.B, C.W, Y.Y, P.P.D, E.Y provided experimental support. S.B.D and I.O helped in CRISPR design and analysis. L.F.A, R.O, O.G, I.G, D.R, I.S and N.K helped with IHC staining and image analysis. All authors contributed to the final version of the paper.

## Supporting information

Supplementary Table 1- WES analysis for all mouse SCCs and UVB-treated samples

Supplementary Table 2- Non-silent mutations per SCC

Supplementary Table 3- Pathway enrichment in samples at day 0 post-inoculation

Supplementary Table 4- Pathway enrichment in samples at days 6, 10, 16, and 20 post-inoculation

Supplementary Table 5- Differentially expressed genes in tumor compartment at days 0, 6, 10, and 16 post-inoculation

Supplementary Table 6- CRISPR pooled guide library constructs

Supplementary Table 7- List of primers used for sgRNA library generation

Supplementary Table 8- List of antibodies used for IHC staining

## Acknowledgments

Y.S. is supported by the Israel Science Foundation grant no. 2133/23, he European Research Council (ERC) under the European Union’s Horizon 2020 research and innovation programme (grant agreement No 770854), by the European Union (ERC, Mel-Immune, 101094980). Views and opinions expressed are however those of the author(s) only and do not necessarily reflect those of the European Union or the European Research Council. Neither the European Union nor the granting authority can be held responsible for them. This research was further supported by the MRA (917324), Minerva (714143), by the Alisa and Peter Savitz Foundation, Les and Cyndy Lederer, Brenda Gruss and Daniel Hirsch, the Donald Gordon Foundation, and the Sigmund and Sofie Englander Foundation, Dwek Institute for Cancer Therapy Research, Weizmann-Brazil Tumor Bank, Laboratory in the name of M.E.H Fund established by Margot and Ernst Hamburger. This research was supported in part by the Intramural Research Program of the National Institutes of Health, NCI. Prof. Samuels is the incumbent of Knell Family Professorial Chair. Prof. Samuels is the Head of EKARD Institute for Cancer Diagnosis Research. A.T.S. was supported by a Career Award for Medical Scientists from the Burroughs Wellcome Fund, a Lloyd J. Old STAR Award from the Cancer Research Institute, a Pew-Stewart Scholars for Cancer Research Award, and the Parker Institute for Cancer Immunotherapy. The images in this paper were acquired at the Advanced Optical Imaging Unit, de Picciotto-Lesser Cell Observatory unit at the Moross Integrated Cancer Center Life Science Core Facilities, Weizmann Institute of Science. This work utilized the computational resources of the NIH HPC Biowulf cluster. (http://hpc.nih.gov). J.A.P. was supported by NIH Training Grant 5T32AI007290.

## Materials and Methods

### Mice

Animals were maintained in a specific pathogen-free (SPF), temperature-controlled (22°C ± 1°C) mouse facility on a reverse 12-hour light, 12-hour dark cycle at the Weizmann Institute of Science. Food and water were given *ad libitum.* Mice were handled using protocols approved by the Weizmann Institute Animal Care Committee (IACUC 05640723-1) in accordance with international guidelines. To generate syngeneic mouse cancer models, 6-week-old female C57BL/6 (purchased from Envigo) and NSG (The Jackson Laboratory) were used.

### Cell line B2905

The murine melanoma B2905 cell line was derived from a UV-irradiated HGF-transgenic mouse in a C57BL/6 background (26). The cell line was grown in RPMI (Biological Industries) containing 10% heat-inactivated FBS (GIBCO), 1% L-glutamine (Biological Industries), 1% Penicillin/Streptomycin antibiotics (Invitrogen) and 12.5mM HEPES (Sigma) buffer. All cells were cultured using standard procedures in a 37°C humidified incubator with 5% CO2. Cells were tested routinely for *Mycoplasma* using a Mycoplasma EZ-PCR test kit (Biological Industries).

### MHC flow cytometry

All SCCs were stained separately for assessment of the expression of MHC class I and II using flow cytometry. Cells were washed twice with PBS, passed through a 70-μm filter (Falcon), and incubated for 30 minutes in MACS buffer (Miltenyi Biotec) in the presence of staining antibodies. Cells were then washed twice, resuspended in MACS buffer, and acquired on CytoFLEX flow cytometer (Beckman Colter). For analysis, Kaluza analysis software (Beckman Coulter) was used.

### Tumor inoculation

For transplantation to C57BL/6 WT mice or NSG immunocompromised mice, 5*10^5^ cells in 100μl PBS were subcutaneously injected into the right lower flank. Tumors were measured using calipers. Tumor volume was calculated using the equation π/6*(smallest diameter)^2^*(largest diameter). Tumors were excised for genomic DNA or RNA purification, at 6, 10, 16, and 20 days post-inoculation. Tumor-derived SCCs that were rejected within the 33 days of monitoring were designated as “rejected”, while the others designated as “non-rejected”.

### Generation of single-cell clones (SCCs)

For SCC generation, cells were plated in 96-well plates, in a 1 cell/well concentration. 10 days after initial plating, cells were monitored and wells that showed more than one focal clone were excluded. Single clones were passaged to establish cell lines. The number of passages was controlled and documented, and low passage cells were used for tumor inoculation.

### T cell suppression assay

Splenocytes were isolated from C57BL/6 mice, enriched for CD8^+^ and CD4^+^ T cells using CD4/CD8 (TIL) MicroBeads (Miltenyi Biotec), and labeled with CFSE (5 M, Biolegend). 1 × 10^5^ cells in 100 μL per well were plated in a 96-well plate pre-coated with anti-mouse CD3ε (1 μg/mL, SouthernBiotech). Tumors were harvested from mice 10 days post inoculation and dissociated using gentleMACS (Miltenyi Biotec). F4/8^+^ macrophages were isolated from these tumors using Anti-F4/80 MicroBeads UltraPure (Miltenyi Biotec), and co-cultured with the stimulated T cells (1 × 10^5^ cells per well) for 48 hours. The dilution of CFSE was evaluated by flow cytometry.

### IFNγ ELISA

To measure IFNγ secretion by the T cells co-cultured with the tumor-derived macrophages in the media of the T cell suppression assay we used Quantikine™ ELISA Mouse IFN-γ Immunoassay (R&D Systems, catalog number #MIF00-1) according to manufacturer instructions. Supernatant samples were taken after 48 hours, and subjected to 1:100 dilution. Each sample was tested in duplicate.

### *Mif* KO using CRISPR/Cas9

Mif expression was abolished in the non-rejected SCCs using CRISPR/Cas9. Two single guides RNA were designed: ‘GACGTCAGACTACGTCCCAA’ and ’AGCCAAGGTGTGCCGGCGGG’. Guide RNAs were chosen using the following tools: the MIT CRISPR design tool (63) and sgRNADesigner (64), in their Benchling implementations (www.benchling.com), SSC (65), and CRISPOR (66). Each guide was cloned into a pSpCaV9(BB)-2A-GFP vector (Addgene, cat# 48138) and transfected into cells of the non-rejected SCCs. GFP+ cells were sorted after two days, each cell seeded in a separate well of a 96-well plate. After two weeks of clonal expansion, the knockout (KO) was confirmed using Sanger sequencing, and SCCs that had WT Mif served as controls for the SCCs that had complete KO of the gene, which was later validated using western blot as well (Abcam, clone ERP12463).

### Single-cell RNA library generation

Tumors were dissociated using tumor dissociation kit and gentleMACS (Miltenyi Biotec) according to the manufacture’s protocol. Single-cell suspensions were hashed using TotalSeq-C anti-mouse Hashtag antibodies (BioLegend) and stained with propidium iodine (#P3566, Invitrogen). Live cells were then sorted, washed, and resuspended with PBS containing 0.04% bovine serum albumin (BSA), and counted using trypan blue staining. Hashed samples were combined and loaded onto the Chromium Controller (10x Genomics) with a targeted cell recovery of 20,000 cells per library. For the non-rejected vs. rejected samples, single-cell gene expression and hash libraries were prepared using the Chromium Single Cell 5’ V(D)J v1.1 Kit with Feature Barcode technology (10x Genomics) according to the manufacturer’s protocol. Samples were sequenced on an Illumina NovaSeq using a 150 (read 1) x 8 (index 1) x 150 (read 2) sequencing configuration. For the SCC35 *Mif* KO samples, single-cell gene expression and hash libraries were prepared using the Chromium Single Cell 5’ v2 Kit Dual Index with Feature Barcode technology (10x Genomics) according to the manufacturer’s protocol. Samples were sequenced using the Illumina NovaSeq system with a 150 (read 1) x 10 (index 1) x 10 (index 2) x 150 (read 2) sequencing configuration.

### sgRNA pool design and cloning

The sgRNA pool was constructed using the previously published protocol for cloning into SCAR vectors (33) (Mkate sgRNA backbone Addgene # 162076, pSCAR Cas9 Blast GFP Addgene # 162074), pLX_EFS_Cre_ppt-del, Addgene # 162073, psPax d64V Addgene # 63586).

Briefly, a sgRNA pool was designed with 4 guides per gene using constructs from the Brie mouse CRISPR Knockout Pooled Library (64). The pool contained 200 non-rejection genes of interest that were significantly upregulated in non-rejected clones compared to rejected clones with adjusted p-value < 0.01 and fold change > 1.5 at any time point (219 genes total; 200 of which were present in the Brie library), 5 non-targeting control guides, as well as 3 depletion controls (*Cd47*, *Adar*, *Cd274*) and 2 enrichment controls (*Pten*, *Ccar1*) selected based on previous literature (32) (**Supplementary Table 5**). The oligonucleotides were annealed and then ligated into pSCAR_sgRNA_puro-mKate-lox2272 that had been digested with the enzyme Bsmb1. Amplification was performed via electroporation and guide representation was confirmed by sequencing.

### Lentivirus production

Lentivirus was generated as previously described (33). Briefly, integrating lentivirus for both Cas9 and sgRNA was generated by overnight transfection of adherent HEK293 cells with either the Cas9 or sgRNA vector and the packaging vectors psPax (Addgene #1226) and pMD2g (Addgene #12259). Integrase-deficient lentivirus was generated by transfecting HEK293 cells with the Cre vector, pMD2g, and a variant of the integrase-deficient psPax-D64 (Addgene #63586). Lipofectamine (Thermo Fisher) was used as the transfection reagent. The virus was collected at 48 and 72 hrs post-transfection, then filtered via a 0.45 uM filtration unit (Millipore). The filtered virus was concentrated using the LentiX concentrator (Takara) at 1500 x g for 45 min. The concentrated supernatant was subsequently aliquoted, flash-frozen, and stored at -80C until use.

### Generation of CRISPR-edited cell lines

Edited SCC lines were generated following the protocol as described by Lane-Reticker et al, 2023 (34). In brief, plated SCCs were infected with pSCAR-Cas9-Blast-GFP lentivirus along with 4 ug/ml of polybrene. The transduction rate was determined by flow analysis of the GFP marker after 48 h. Once the transduction rate was confirmed to be sufficient, edited cells were selected with blasticidin for 6 days or until Cas9- expressing cells were greater than 90% of the population. Cas9-expressing cells were then infected with lentivirus for the pSCAR library and underwent selection with puromycin after 48 h. Cells were selected for 10 days or until edited cells were greater than or equal to 90% of the population. After 10 days, cells were infected twice with IDLV-Cre lentivirus in media with polybrene. Using the same process as the other lentivirus infections, cells were monitored for 10 days via flow cytometry for the loss of mKate2 and GFP. After 10 days, greater than 90% of the population was double-negative for fluorescent reporters.

### Tumor inoculation for *in vivo* CRISPR experiments

CRISPR-edited SCC cells were collected and then washed in PBS. 0.5 million cells in 100 µl phosphate buffered saline (PBS) were injected subcutaneously into the left flank of C57BL/6 mouse (one tumor per mouse) and measured every 3 days starting 6 days post-injection. 20 days after injection, tumors were collected, weighed, and then minced into small pieces. The minced tumors were dissociated using the Miltenyi tumor dissociation kit and a gentleMacs octo dissociator with the soft/medium tumor program, then subjected to red blood cell (RBC) lysis. The resulting cell suspension was then passed through a 70 µm filter and incubated in FACS buffer (PBS supplemented with 1% BSA and 2mM EDTA) in the presence of Fc block and staining antibody. Live CD45-tumor cells were then isolated by FACS before proceeding to library preparation and sequencing.

### sgRNA library preparation and sequencing

Genomic DNA was extracted using a commercially available kit (Zymo cat# D3025). SgRNA libraries were prepared for sequencing as previously described (67). Briefly, a standard three-step amplification was used using the P5 and P7 primers listed in (**Supplementary Table 7**). First, sgRNAs were amplified from gDNA in 100 µl reactions with up to 4 µg of gDNA used per reaction for 22 cycles. For sequencing of plasmid pools, this first PCR was skipped. For the second PCR, 0-7bp offset was added to the library using pooled stagger primers to increase the diversity of the library, with PCR 2 primers targeting sites nested inside of PCR 1 products. Finally, libraries were indexed and then sequenced in dual-indexed 1x75 bp format on an Illumina NextSeq.

### WES and RNAseq data processing

Genomic DNA was extracted from cell lines of the SCCs using the QIAGEN DNeasy Blood & Tissue Kit. Exome capture was performed using the SureSelectXT Mouse All Exon System (Agilent Technologies). Mapping of WES FASTQ reads to the mouse genome (GRCm38, mm10) was implemented using BWA mem-O 6 -E 1 -B 4 -t 4 -M -R “@RG ID:bwa LB:1 SM:s PL:illumina” Mus_musculus.GRCm38.all.fa S_R1.fastq.gz S_R2.fastq.gz (68). Sorting and indexing of the BAM files was performed using samtools v1.8 (69). Subsequently, the Picard MarkDuplicates module was applied. Samtools mpileup(69) and bcftools v1.3 (69) were used for variant calling. Mouse SNP filtering was applied to the VCF files based on normal mouse spleen and kidney SNPs derived from the injected mouse strain. Furthermore, SNPs obtained from the B2905 parental cell line were removed. RNA was extracted from cell lines and tumors of the rejected and non-rejected SCCs at days 0, 6, 10, 16 and 20 post-inoculation, followed by bulk MARS-seq library preparation (70). Sequencing was done using Illumina NextSeq 500 sequencer using the Nextseq High Output V2 150 sequencing kit (Illumina, 150 bp, single reads). RNAseq bulk sequencing data (from MARS-seq) was analyzed using the UTAP trancriptome pipeline tool (71).

### Principal component analysis (PCA) of bulk gene expression data

The RNA-seq read count data were TMM-normalized and log-CPM-transformed with the edgeR package (72) and then subjected to a principal components analysis (PCA) applying variable standardization. The samples were visualized on a PCA plot showing the top 2 principal components, and after visual inspection, several outlier samples were identified (including “scc35day16c”, “scc31day6c”, “scc31day10c”, “scc40day10c”, “scc32day20a”, “scc40day6a”, “scc32day6a”, “scc37day6c”, “scc35day6b”, and “scc35day0b”) and were excluded from the final PCA plot and further analysis. These identified outliers have potential quality issues with higher Ct values and are from the same pool of RNA library preparation.

### Analysis of somatic mutation data

VEP (73) was used to annotate the called variants with the reference genome GRCm38 from Ensembl. From the VEP annotation, we extracted information on the gene transcripts mapped to each mutation, as well as the effect of the mutation on the protein level (e.g. animo acid changes, if any). Non-silent mutations, as described in the main text, were defined as a DNA-level variant with an annotated amino acid change or annotated as a splicing site variant in any of its associated transcripts.

### Variant Allele Frequency

For each somatic mutation that survived the above-described filtering, the Variant Allele Frequency (VAF) was calculated. This is defined by the number of reads in support of the somatic mutation divided by the total number of reads covering the position of the mutation. The distribution of VAFs was plotted using the probability density function. A threshold of VAF=0.25 was set to separate between clonal and subclonal mutations similar to the definition of Williams et al. 2016 (74).

### Differential expression (DE) and pathway enrichment analysis of bulk gene expression data

DE analysis comparing the rejected and non-rejected clones was performed with the limma-voom method (75). The DE test was made at each time point, and we also tested for the average DE across all *in vivo* time points (i.e. after transplantation); the tests were conducted using nested design models taking into consideration replicated samples of each clone. We note that we did not use edgeR for testing (although we used edgeR’s TMM normalization to obtain a normalized expression matrix as input to several other analyses, e.g. PCA) because it was not able to handle the more flexible nested design model we used for across *in vivo* time points. GSEA (76), implemented in the fgsea package (77), was used for pathway enrichment analysis based on DE log fold-change values, with pathway definitions taken from the Reactome database (78). The Benjamini-Hochberg method was used for multiple hypothesis correction of P values (79). For visualizing the pathway enrichment more intuitively, GSVA (80) was used to compute per-sample pathway activity score (i.e., the GSVA enrichment score) using the TMM-normalized log-CPM expression values, and these scores were used in various plots (but they were not directly used for statistical testing).

### Phylogenetic analysis of mouse UVB and SCCs

WES data for the UVB and the 40 SCCs were used for joint clustering to infer the subclones present across this combined set of samples. Copy number alterations were utilized as input to the SciClone algorithm (81). The following filters were applied: a minimum alternative read depth of >5, indels and triallelic sites were excluded, and only variants present in ≥2 samples were retained. The running parameters were: copyNumberMargins=0.5, maximumClusters=30, minimumDepth=-1. A phylogenetic tree was constructed using the R package CloneEvol (82).

### Differential expression of rejected vs. non-rejected tumor clones

To characterize differences between rejected vs. non-rejected SCCs, differential expression analysis was performed on tumor cells at each time point (day 0, 6, 10, and 16 post-inoculation) separately using the Wilcoxon rank sum test. Significant differentially expressed genes were defined as those with a 1.5-fold difference in mean log-normalized expression between the rejected vs. non-rejected group and a Bonferroni-corrected adjusted p-value < 0.01. Core rejection and non-rejection signatures were defined per time point (day 0, 6, and 10 post-inoculation) by averaging the expression of significant genes per SCC, scaling the averaged expression values across all SCCs, and identifying genes that displayed a consistent pattern among all rejected or non-rejected SCCs respectively (i.e., the core non-rejected signature includes genes for which z-score > 0.1 in all non-rejected SCCs and < 0 for all rejected SCCs). The core signature analysis was not performed for the day 16 time point since there was only one rejected clone (SCC40) present.

### Estimating cytolytic activity from RNA-seq data

Gene expression levels (RPKM) of the 330 melanoma patients with corresponding survival information were downloaded from the TCGA portal (83). Cytolytic activity (CYT) of TILs in patient tumors was estimated from the geometric mean of expression levels of GZMA and PRF1.

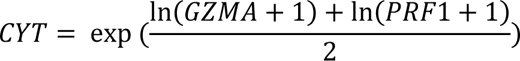

### Computation and analysis of T cell function and T cell infiltration scores using bulk gene expression data

The TIDE (27) algorithm was applied to the bulk RNA-seq data to compute measures of T cell function from gene expression. We used the offline TIDEpy package with default normalization and parameters. TIDE was initially used to predict immune checkpoint blockade (ICB) therapy response via computing two different scores for each sample: a T cell “dysfunction” score and a T cell “exclusion” score. In principle, these scores can be used to reflect anti-tumor immune function and we used TIDE for this alternative purpose. The T cell infiltration score we described in this study is the negative of the “exclusion” score from TIDE. The “dysfunction” score from TIDE is higher in ICB responders and thus appears to be a misnomer since this score positively correlates with T cell function. Therefore, we call it the T cell function score in this study. The differences in these scores between the rejected and non-rejected clones were tested with a linear model at each time point; the tests were conducted using nested design models taking into consideration replicated samples of each clone. The Benjamini-Hochberg method was used for multiple hypothesis correction of P values.

### T cell exhaustion score analysis

A T cell exhaustion signature (a plastic dysfunctional state from which T cells can be rescued) of 25 genes from Philip et al. 2017 (84) was used to score the exhaustion of individual CD8+ T cells in each clone.

### Data processing of hashed scRNA-seq libraries

Reads from scRNA expression libraries were aligned to mouse genome assembly GRCm38 (mm10) and quantified using cellranger count (10x Genomics, version 6.0.0). The filtered feature-barcode matrices containing only cellular barcodes were used for further analysis. Single-cell gene expression matrices were imported into R (version 4.2.0) and analyzed using Seurat (version 4.3.0) (85). Cells with number of genes captured between +/- 2 standard deviations of the mean per library were kept. Additionally, cells with greater than 10% mitochondrial RNA reads were excluded from subsequent analyses. Libraries were demultiplexed using HTODemux() with a positive quantile of 0.99. Only cells that were identified as a singlet were kept for downstream processing. For each library, log normalization and variable feature selection based on variance stabilizing transformation were first performed on each sample individually. Then, anchors between samples were identified using FindIntegrationAnchors() and integrated using IntegrateData() with 30 dimensions. The integrated expression matrix was then scaled and centered for each feature. Next, linear dimensional reduction was performed on the integrated scaled data using PCA. For visualization, uniform manifold approximation and projections (UMAPs) were calculated using the first 30 principal components. To identify clusters based on gene expression profiles, we performed shared nearest neighbour (SNN) modularity optimization clustering.

### Calculation of M2 macrophage score

Cell cluster/type compositions are plotted using the dittoBarPlot() function from the dittoSeq package (86). The annotation of M1 macrophages and M2 macrophages is based on the Tumor Immune Single-Cell Hub (TISCH) (39) annotation for the BRCA_GSE114727_inDrop dataset published by Azizi et al. 2018 (87). The top 100 marker genes for each cell type were downloaded from TISCH and used to calculate the M1 and M2 score for each cell using the Seurat AddModuleScore() function. Violin plots of gene signature scores are calculated using the Seurat VlnPlot() function.

### Gene expression patterns analysis

Expression patterns of the C1q subunit and *Apoe* genes are plotted using the Seurat FeaturePlot() function.

### Opal multiplexed IHC staining

For Opal multiplexed immunohistochemistry (IHC) staining, tumors were excised at days 10 post-inoculation, fixed in 4% (w/v) paraformaldehyde for 24 hours, and restored in 1% paraformaldehyde until embedded in paraffin for histological analysis. Immunohistochemistry was performed on deparaffinized and rehydrated 4-μm thick paraffin-embedded sections using xylene and a decreasing concentration of Ethanol (100%, 96%, 70%). Endogenous peroxidase activity was blocked with 3% H2O2 and 1% HCL in Methanol for 30 min, followed by heat-induced antigen retrieval in Tris-EDTA (PH=9). For nonspecific binding, sections were blocked with 20% NHS (Vector Labs -S-2000) and 0.1% triton. In case a secondary biotinylated antibody was used, an additional step of biotin blocking kit (Vector Labs -SP-2001) was performed. Primary antibodies, diluted in 2% NHS and 0.1% triton, were incubated overnight according to the **Supplementary Table 8**. Secondary HRP or biotin-conjugated antibodies incubation was followed by fluorescently labeled OPAL reagents or streptavidin. CD3 staining was double amplified using a biotinylated secondary antibody, followed by ABC kit (Vector Labs -PK-6100) and OPAL reagent. Antibodies were removed using 10 minutes of microwave treatment with Tris-EDTA (pH=9), and then the protocol repeated from the blocking step. Nuclei were stained with DAPI (not shown).

### Multiplex fluorescence imaging

Multispectral imaging was performed using PhenoImager at 20× magnification (Akoya Biosciences, Marlborough, MA) according to manufacturer instructions. Later, the multispectral acquired images were loaded into InForm software for unmixing and background subtraction (inForm v.3.0; Akoya Biosciences). In the inForm software, an unstained slide (without DAPI and OPAL staining) was loaded and regions with high autofluorescence signals were marked for processing. After processing, image tiles were stitched using QuPath software with the “merge multiple TIFF fields” script.

### Image processing

To detect and quantify the macrophages (F4/80^+^ CD204^+^ CD206^+^) and T cells (CD3^+^ CD8^+^), cell segmentation and classification was applied to the fluorescent-markers stained images using QuPath (v0.4.3) (88). Nuclei were segmented from the 4’,6-diamidino-2-phenylindole (DAPI) channel using StarDist (89) and further inflated to have approximated cell segmentation. The cells were then classified as positive or negative for each of stains, and the total number of double or triple positive cells and their respective ratio were quantified within 5 representative regions of 500*500um for each tumor. To improve nuclei segmentation a new StarDist model was trained using the ZeroCostDL4Mic (90) StarDist notebook with examples that were not perfectly segmented by the provided model. Positive/Negative random trees cell classifiers were trained for each stain independently on multiple image regions representative of the tissue characteristics and experimental conditions. The classifiers were then combined together to detect double and triple positive cells. The classifiers were then applied to selected representative areas of the tumors, based on a threshold classifier, to quantify the total number of the double (CD3^+^ CD8^+^ T cells) and triple (F4/80^+^ CD204^+^ CD206^+^ macrophages) positive cells, and their respective ratio to the total number of cells per tissue regions of interest.

### CRISPR pooled screen data analysis

sgRNA libraries from the pooled CRISPR screens were analyzed using the MAGeCK pipeline (91). Briefly, sgRNA counts are first computed per library using mageck count and median-normalized in order to adjust for library sizes and read count distributions. Then, sgRNA libraries sequenced at 20 days post tumor inoculation were compared to the input guide library using the mageck test to identify enriched and depleted guides and genes.

### Quantification and Statistical Analysis

Statistical analysis was performed using the Prism 10 software (GraphPad, San Diego, CA, USA) and the software environment R, using RStudio. All data is presented using standard error mean (SEM). P values are depicted in all figures, and selected p values with exceptional significance to the paper are also briefly described in the main text. Samples sizes (n), means and SEM are depicted in the figures and/or figure legends. Sample size values were either depictions of the number of mice used for experiments or the number of patients.

### TCGA survival analysis

To observe the effect of *MIF* expression in tumor cells on overall patient survival, we stratified the TCGA (469 melanoma) patients into high (≥ median) and low (< median) *MIF* expression in deconvolved gene expression of TCGA data (37). We further checked for the combination of Tumor Infiltrated Lymphocyte (TIL) patterns (38) within high/low *MIF* expression for 377 TCGA melanoma patients and observed the effect on overall patient survival. We fit Kaplan-Meier survival curves for each group to test for any significant survival differences between the groups using the Log-rank test. R packages ‘survival’ and ‘survminer’ were used to perform all the survival analyses (92).

To test for an association between ITH and overall patient survival, we considered 262 TCGA melanoma patients without censoring information. We used the previously established calculation of ITH based on Wolf et al. 2019 (5).

### Data availability

All scRNA-seq datasets generated in this study have been deposited in NCBI’s Gene Expression Omnibus (GEO) and are publicly accessible under accession number GSE247059.

## Table legends

**Supplementary Table 1- WES analysis for all mouse SCCs and UVB-treated samples.** The number of mutations and extent of clonality are calculated for each sample. The proportion of C>T (and complementary G>A) alterations is provided as expected following UVB exposure in melanoma.

**Supplementary Table 2- Non-silent mutations per SCC.** List of the non-silent mutations in each SCC, who passed all the filtration steps of WES data processing described in the methods. Supplementary table to **Supplementary Figure 1**E.

**Supplementary Table 3- Pathway enrichment in samples at day 0 post-inoculation.** Significantly enriched pathways comparing rejected clones to non-rejected clones. Direction is indicated by the sign in the “NES” (normalized enrichment score) column. A positive NES has higher expression in the rejected compared to the non-rejected clones.

**Supplementary Table 4- Pathway enrichment in samples at days 6, 10, 16, and 20 post-inoculation.** Averaged effect across all *in vivo* time points. Significantly enriched pathways comparing rejected clones to non-rejected clones. Direction is indicated by the sign of the “NES” (normalized enrichment score) column. A positive NES has higher expression in the rejected compared to the non-rejected clones.

**Supplementary Table 5- Differentially expressed genes in tumor compartment at days 0, 6, 10, and 16 post-inoculation.** Differential expression analysis between tumor cells from rejected and non-rejected clones. Significance was assessed by Wilcoxon rank sum test and p-values were adjusted using Bonferroni correction. The ‘avg_log2FC’ column represents the fold change between the average gene expression among rejected SCCs versus the average gene expression among non-rejected SCCs. Whether a gene is overexpressed by all clones within the rejected or non-rejected categories is indicated in the ‘core_genesig’ column.

**Supplementary Table 6- CRISPR pooled guide library constructs.** sgRNA library for the pooled *in vivo* screen targeting non-rejection genes. sgRNA sequences are taken from the Brie mouse CRISPR Knockout Pooled Library (64).

**Supplementary Table 7- List of primers used for sgRNA library generation.** P5 and P7 primers used to amplify and sequence the sgRNA library pool for the non-rejection CRISPR screen.

**Supplementary Table 8- List of antibodies used for IHC staining.** List of primary and secondary antibodies used for the Opal staining, as well as staining conditions and Opal fluorophores.

---

**Supplementary Figure 1.**
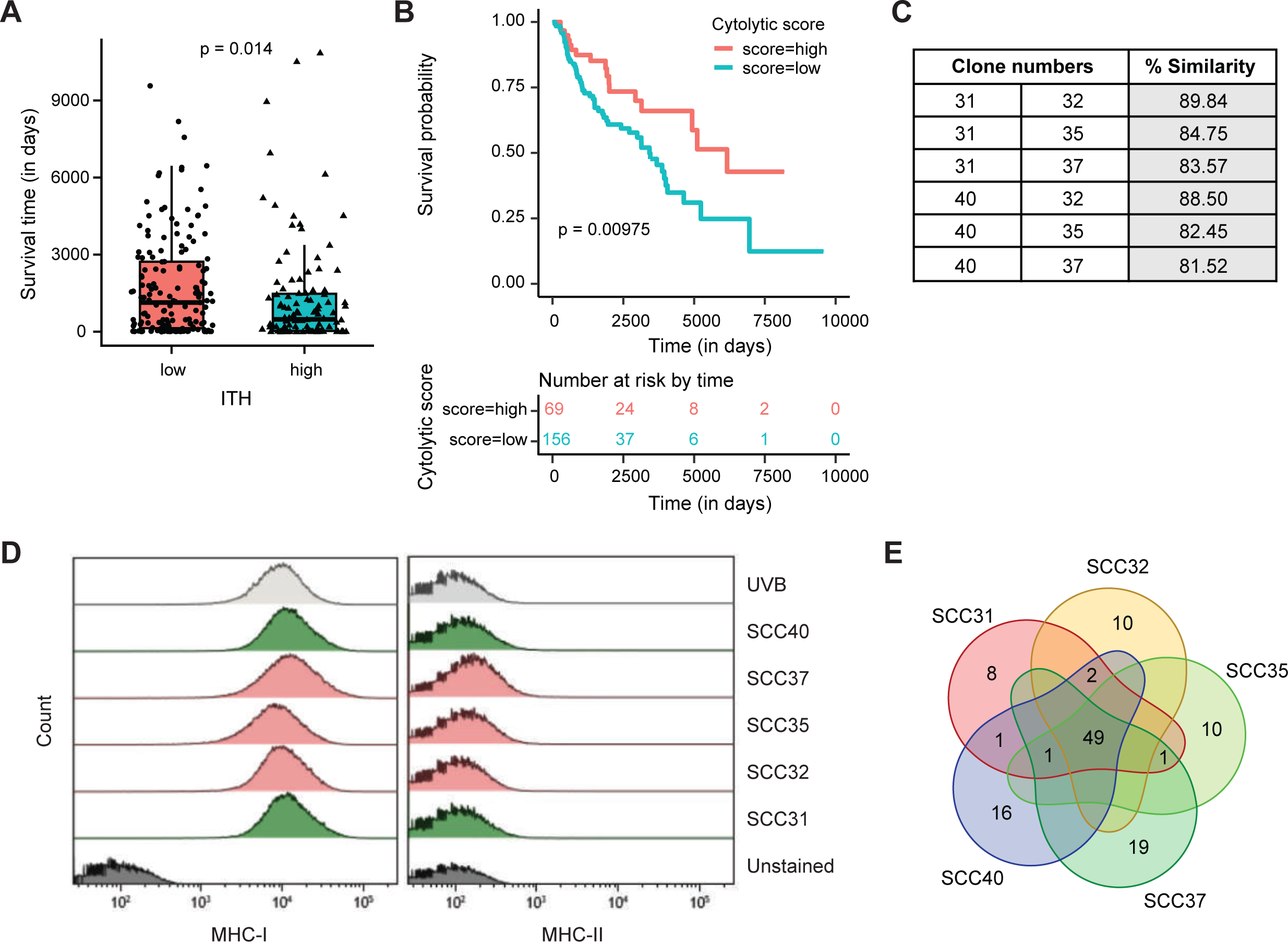
Clinical data and characterization of the Single-cell clones. A) Boxplot of survival times (in days) for low vs. high ITH patients for TCGA melanoma patients (excluding censored data). The individual points denote the survival time for patients in the respective categories. P-value was determined using the Wilcoxon’s test between low and high ITH patients, without censoring. **B)** Kaplan-Meier survival curves (time is measured in days on the x-axis) of the cytolytic score of TCGA melanoma patients with low ITH. P-values were determined using the log-rank test to compare the overall survival probability between patients with high (in red) and low (in blue) cytolytic scores. Patients with high cytolytic score have significantly better survival probabilities. **C)** Genetic similarity percentage for each pair of rejected and non-rejected clones. **D)** Evaluation of MHC-I and MHC-II levels in each SCC, measured by flow cytometry. Green color represents the rejected SCCs, pink represents the non-rejected SCCs, and grey represents the parental UVB irradiated cell line. **E)** Somatic mutation profiles of the rejected and non-rejected SCCs. A Venn diagram showing the overlaps of non-silent somatic mutations across the clones, based on whole exome sequencing data. The numbers of non-silent somatic mutations shared and unique to the clones are labeled in the diagram, regions without labels contain no non-silent somatic mutation. The SCC31 (red) and SCC40 (blue) clones on the left-hand side are rejected, the rest are non-rejected.

**Supplementary Figure 2.**
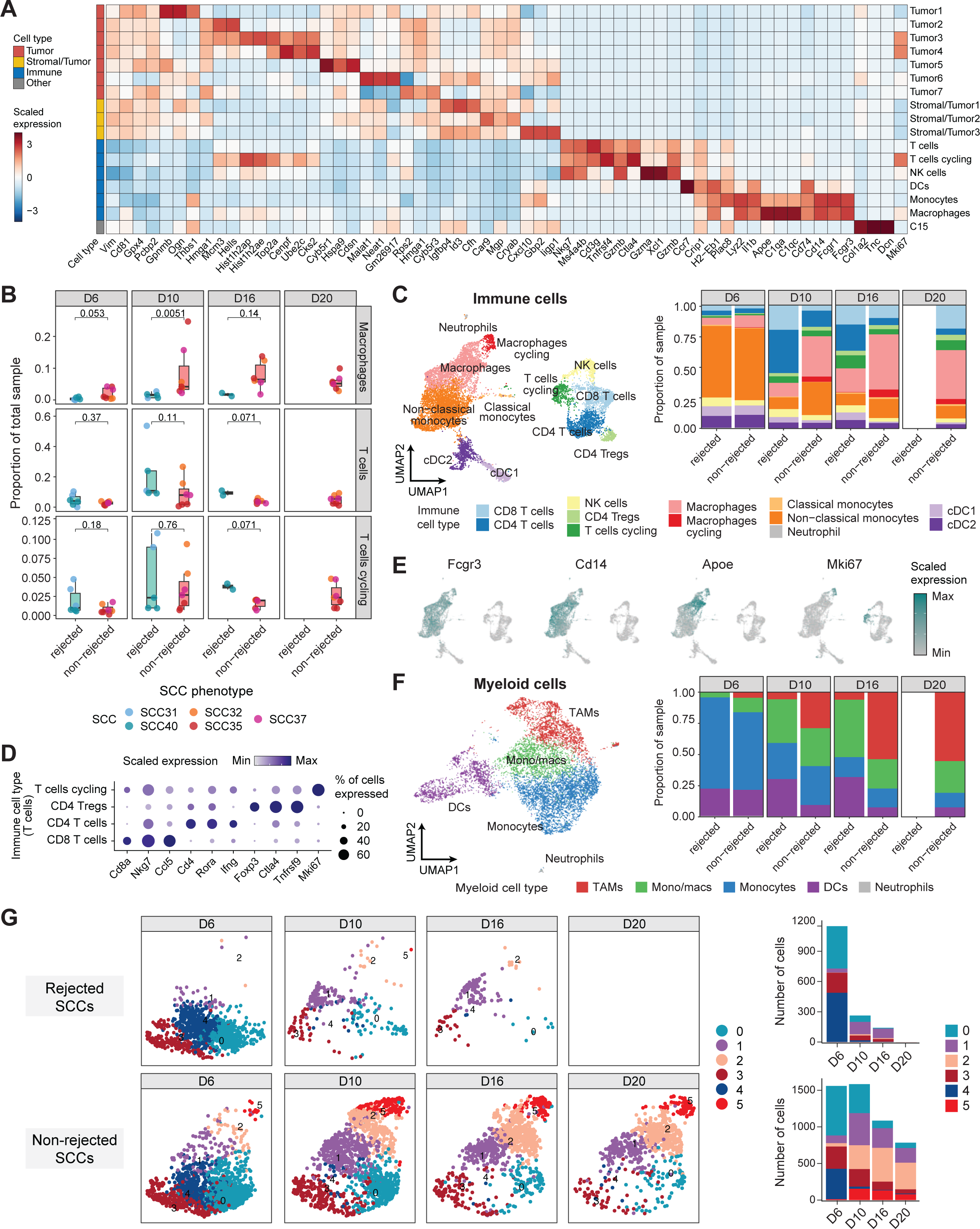
Cell types from scRNA-seq of tumors from mice inoculated with rejected or non-rejected SCCs. A) Heatmap of cluster marker expression among cell type clusters derived from scRNA-seq of rejected and non-rejected SCCs. Color indicates z-score normalized expression per gene. **B)** Boxplot of cell type proportions of macrophage (top) and T cell (bottom) clusters from rejected and non-rejected SCCs at day 6, 10, 16, and 20 after tumor inoculation. Statistical testing by two-sided Wilcoxon test. **C)** UMAP of re-clustered immune cell clusters obtained from scRNA-seq of tumors from mice inoculated with rejected or non-rejected SCCs (left). Bar plots of immune cell type proportions per SCC group at day 6, 10, 16, and 20 after tumor inoculation (right). **D**) Dot plot showing expression of T cell genes among the T cell clusters presented in (**C**). **E)** UMAP expression plots of *Fcgr3* and *Ki67* among cell-type clusters of the immune compartment. **F)** UMAP of re-clustered myeloid cell clusters obtained from scRNA-seq of tumors from mice inoculated with rejected or non-rejected SCCs (left). Bar plots of myeloid cell type proportions per SCC group at day 6, 10, 16, and 20 after tumor inoculation (right). **G)** Distribution and composition of Mon/Mac subclusters between different time points in rejected (top) and non-rejected (bottom) groups.

**Supplementary Figure 3.**
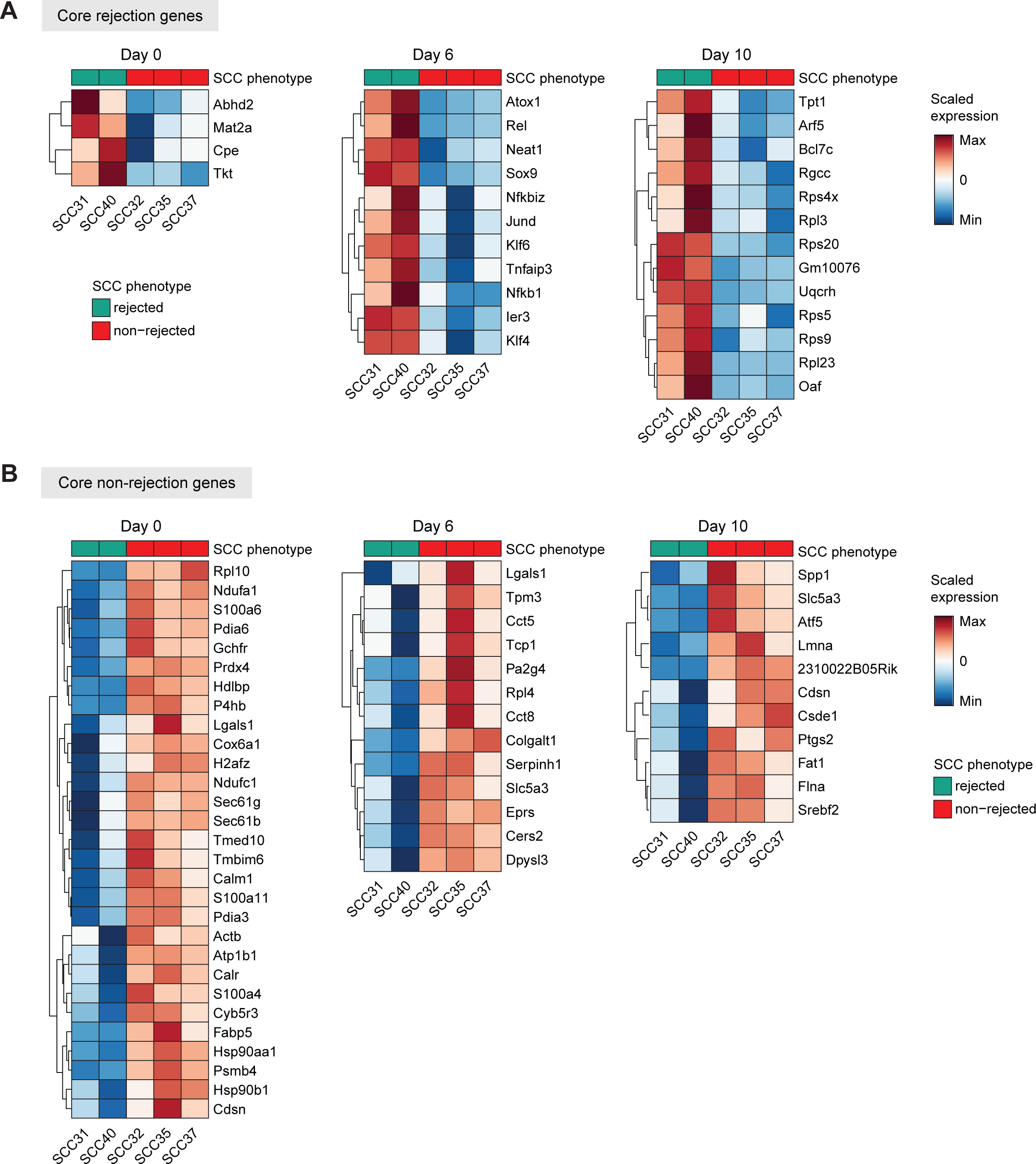
Differential expression analysis of rejected vs. non-rejected tumor cells. A- **B)** Differentially expressed genes in the rejected **(A)** vs. non-rejected **(B)** SCCs at different time points post-inoculation.

**Supplementary Figure 4.**
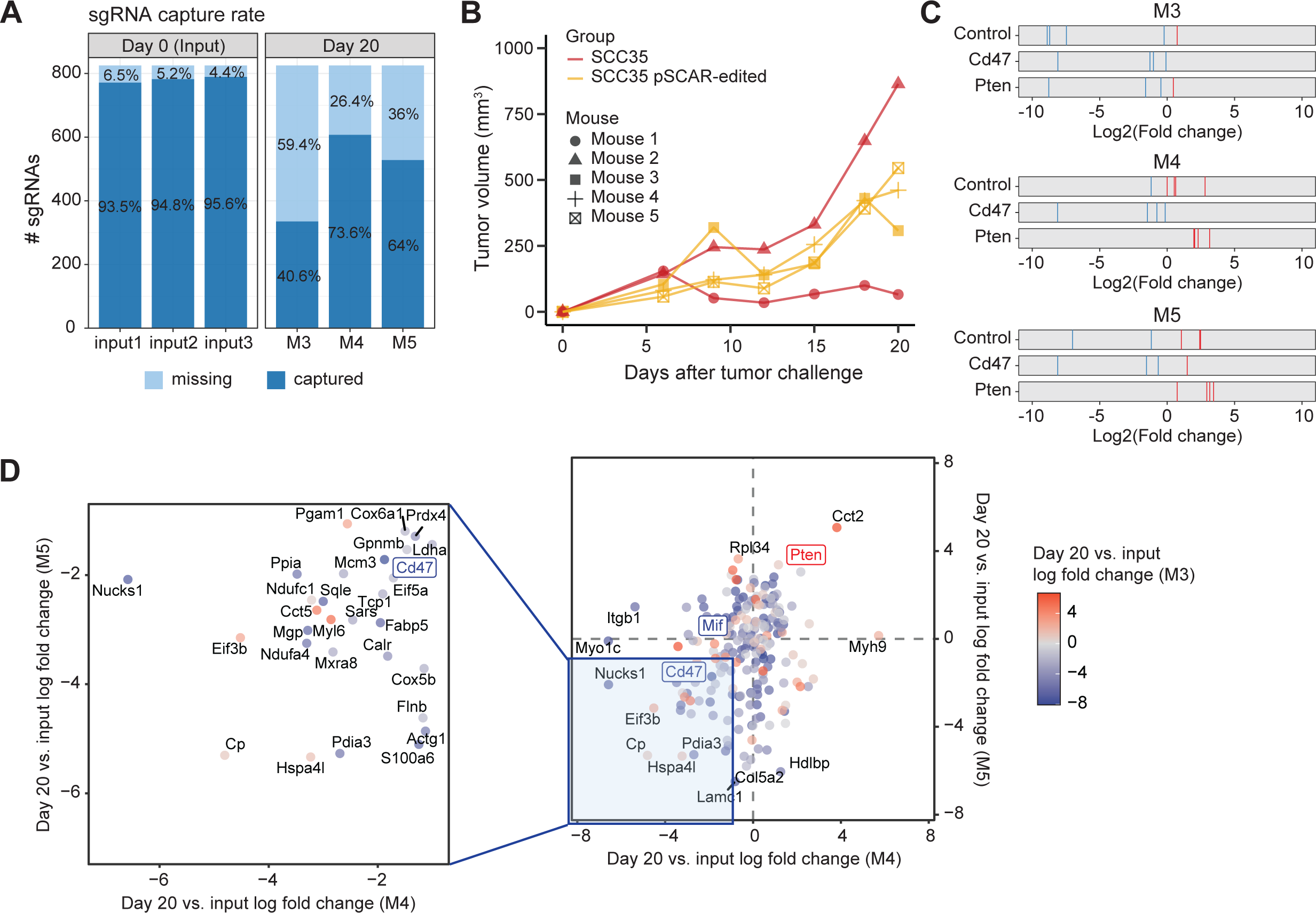
Supplementary data for the *in vivo* CRISPR screen. A) Capture rate of the sgRNA pool among CRISPR-edited SCC35 libraries. Numbers indicate the percentage of sgRNAs that are missing or captured in each library. **B)** Growth curves of SCC35 (n=2) and CRISPR-edited SCC35 (n=3) tumors in immunocompetent mice. **C)** Log fold change of non-targeting control guides and sgRNAs targeting *Cd47* (depletion control) and *Pten* (enrichment control) between day 20 post-inoculation and day 0 input. Enriched and depleted guides are colored with red and blue lines respectively. **D)** Gene-level depletion and enrichment of CRISPR-edited libraries. The score represents the average log2-fold change of guide counts at day 20 post-inoculation and day 0 input libraries. Highlighted in the left square are the top depleted genes ranked by the score.

**Supplementary Figure 5.**
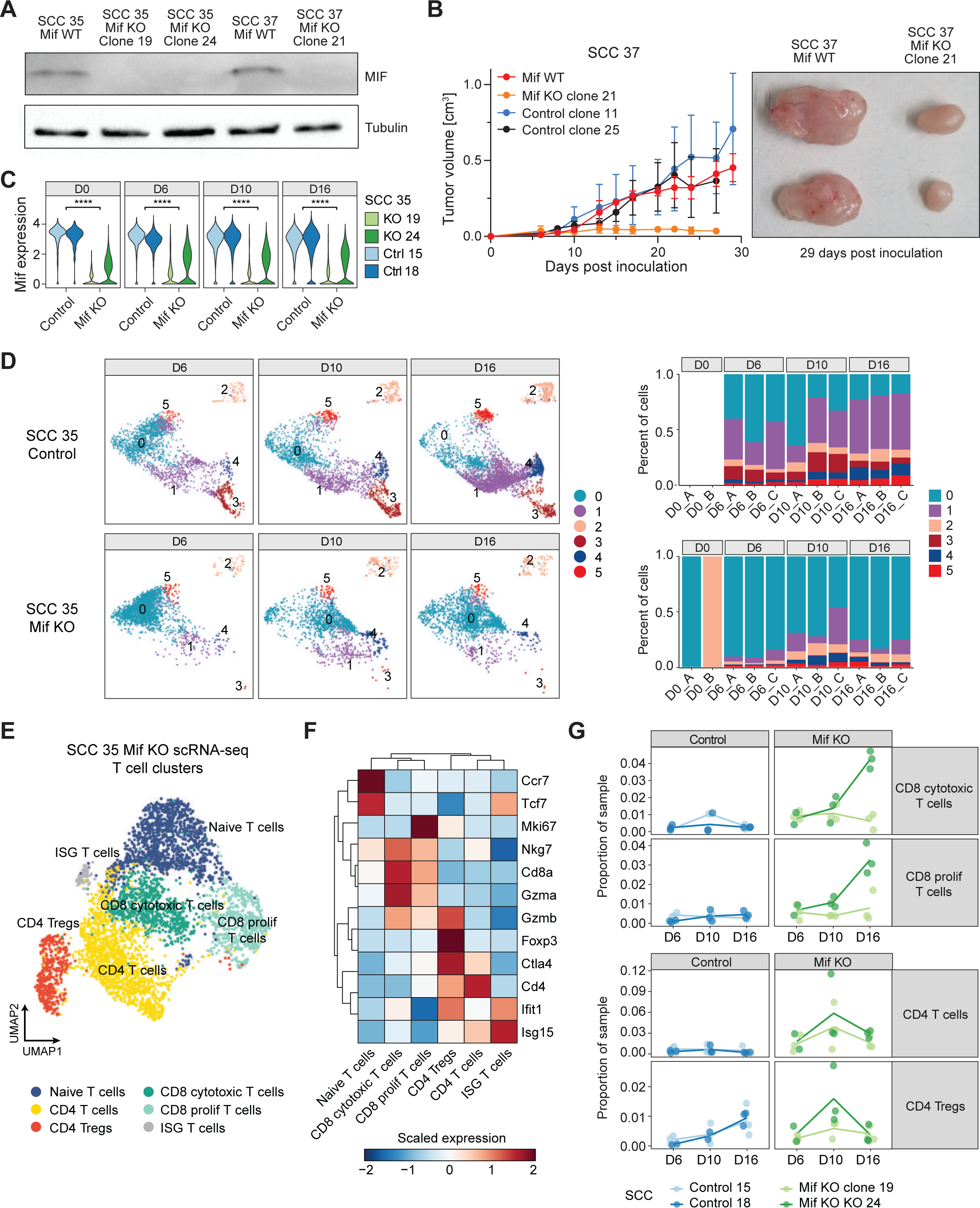
scRNA-seq characterization of *Mif* KO tumor clones. A) Western blot for MIF and Tubulin in SCC35 and SCC37 cells. **B)** *Mif* KO using CRISPR in the non-rejected SCC37 significantly decreased tumor growth compared to the *Mif* WT control clones. Statistical analysis between *Mif* KO to WT expressing clones at day 27 (the latest time point in which all groups are represented) gave a P value of 0.00004 by two-sided Wilcoxon-test followed by Bonferroni correction. n=7-18 **C)** Violin plots of *Mif* expression among tumor cells derived from scRNA-seq of *Mif* KO and control tumors at days 0, 6, 10, and 16 post-inoculation. Statistical testing by two-sided Wilcoxon-test (**** indicates p-value < 0.0001). **D)** Distribution and composition of Mon/Mac subclusters between different time points in *Mif* KO (top) and control (bottom) groups. **E)** UMAP of re-clustered T cell populations from SCC35-derived tumors with or without *Mif* KO. **F)** Heatmap of T cell marker expression among T cell clusters from scRNA-seq of *Mif* KO and control tumors. Color indicates z-score normalized expression per gene. **G)** Longitudinal frequencies of re-clustered CD8 (top) and CD4 (bottom) T cell types from **(E)** in control and *Mif* KO tumors.

